## Supplementary material for "Somatostatin-expressing neurons regulate sleep deprivation and recovery": Figures_S42-S44.pdf

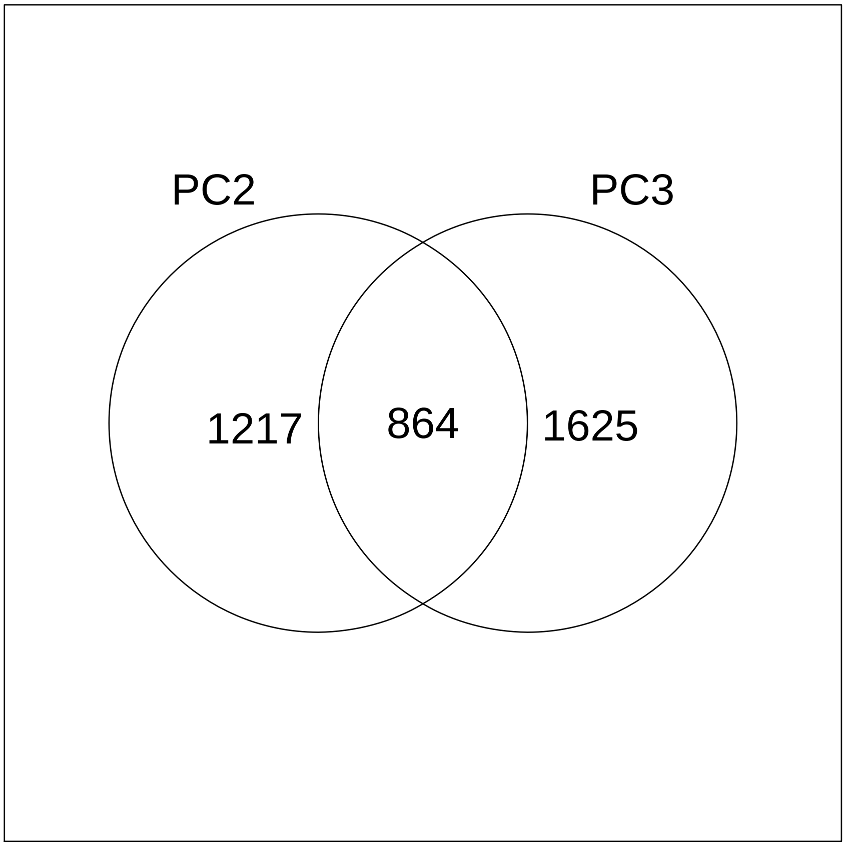


Figure S42: Venn diagram between gene symbols selected by $\ell_1 = 2$ and $\ell_1 = 3$, respectively. PC2: $\ell_1 = 2$ and PC3: $\ell_1 = 3$.

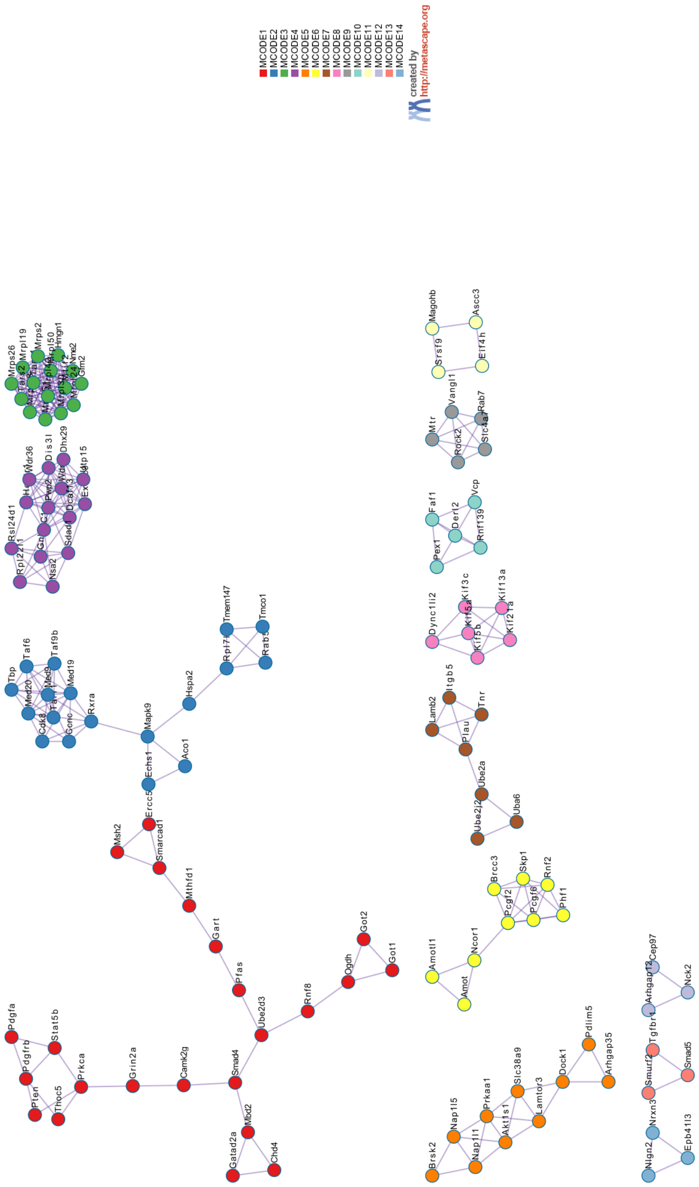


Figure S43: Protein–protein interaction network and MCODE components identified in the gene lists. For annotation of cluster, see Tables ?? and ??.

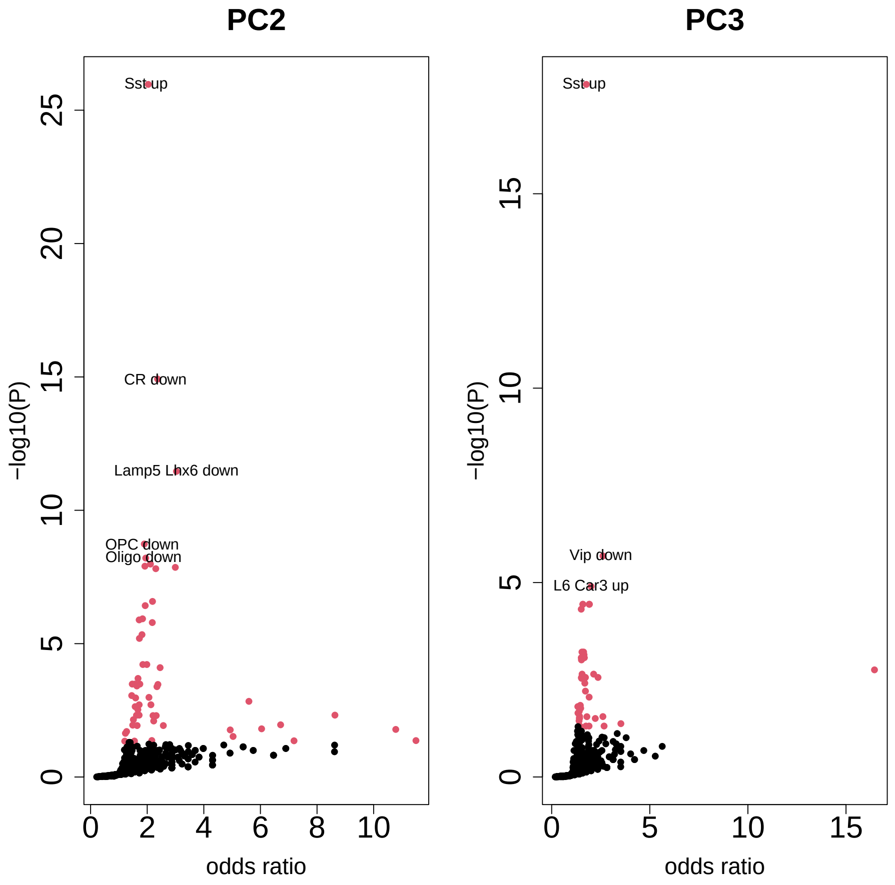


Figure S44: Volcano plot of cell clusters in "Allen Brain Atlas 10x scRNA 2021" category. Red ones are associated with adjusted $P$-values less than 0.05. PC2: $\ell_1 = 2$ and PC3: $\ell_1 = 3$.
