## Supplementary material for "Somatostatin-expressing neurons regulate sleep deprivation and recovery": README.txt

Supplementary file 1: Figure S1 Full list of enrichments using Metascape (Figure_S1.pdf)

Supplementary File 2: ZIP files as described above (Metascape)

Supplementary file 3: Selected genes and Allen Brain Atlas for $\ell_1=2$ (genes_PC2.ods)

Supplementary file 4: Selected genes and Allen Brain Atlas for $\ell_1=3$ (genes_PC3.ods)

Supplementary file 5: Supplementary Tables S1–-S20 in Table 6 (Supplementary_Tables.pdf)

Supplementary file 6: Supplementary Figures S2–-S21 in Table 6  (Supplementary_Figures.pdf)

Supplementary file 7: Supplementary Figures S42–-S44 (Figures_S42-S44.pdf)
