## Supplementary material for "Somatostatin-expressing neurons regulate sleep deprivation and recovery": Supplemetrary_Figures.pdf

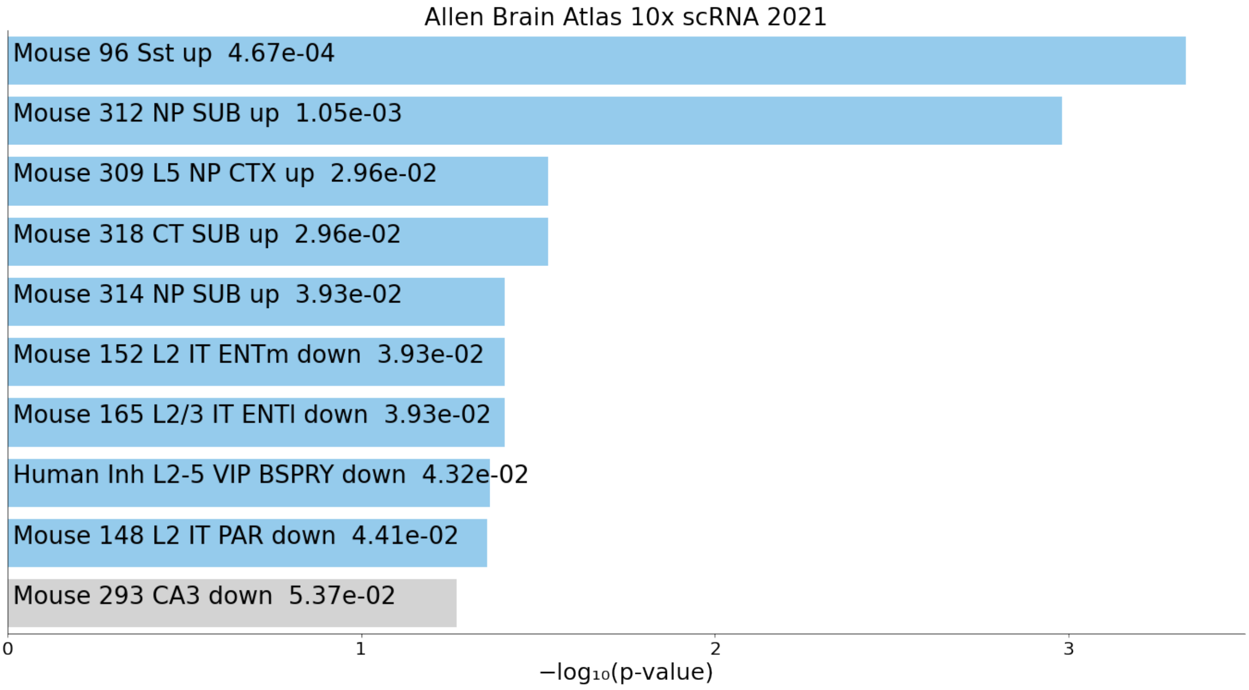


Figure S2 left_cerebral_cortex_up1

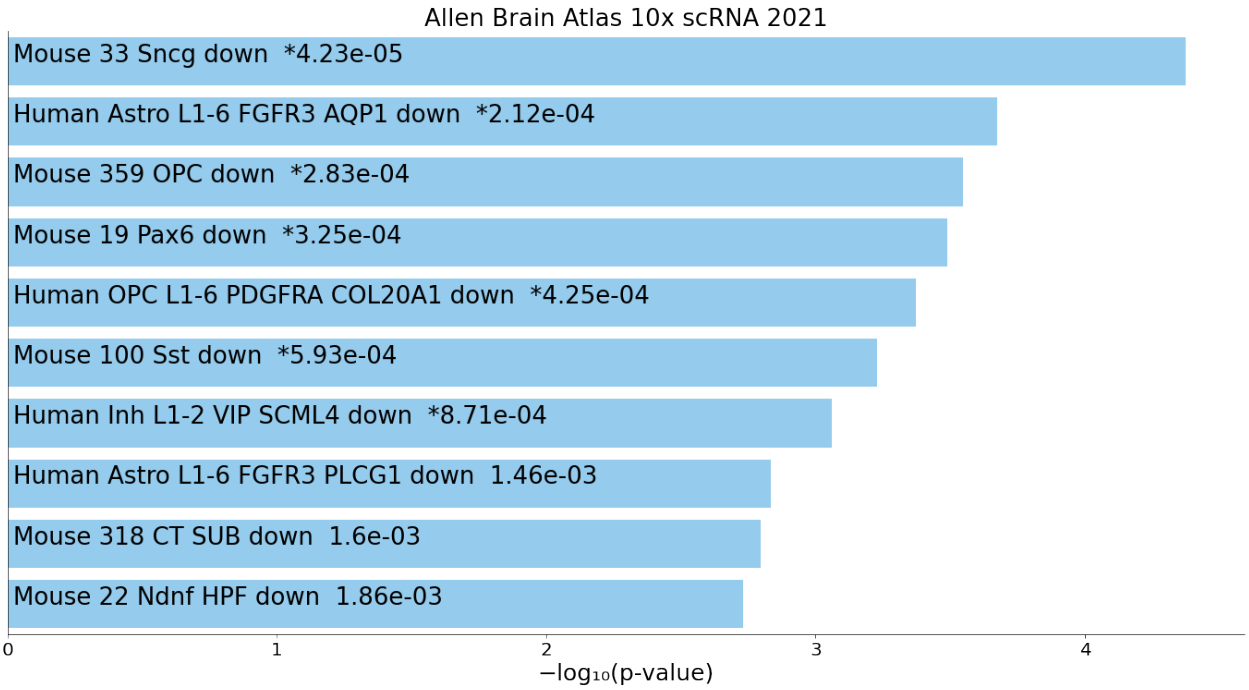


Figure S3 left_cerebral_cortex_down1(brain)

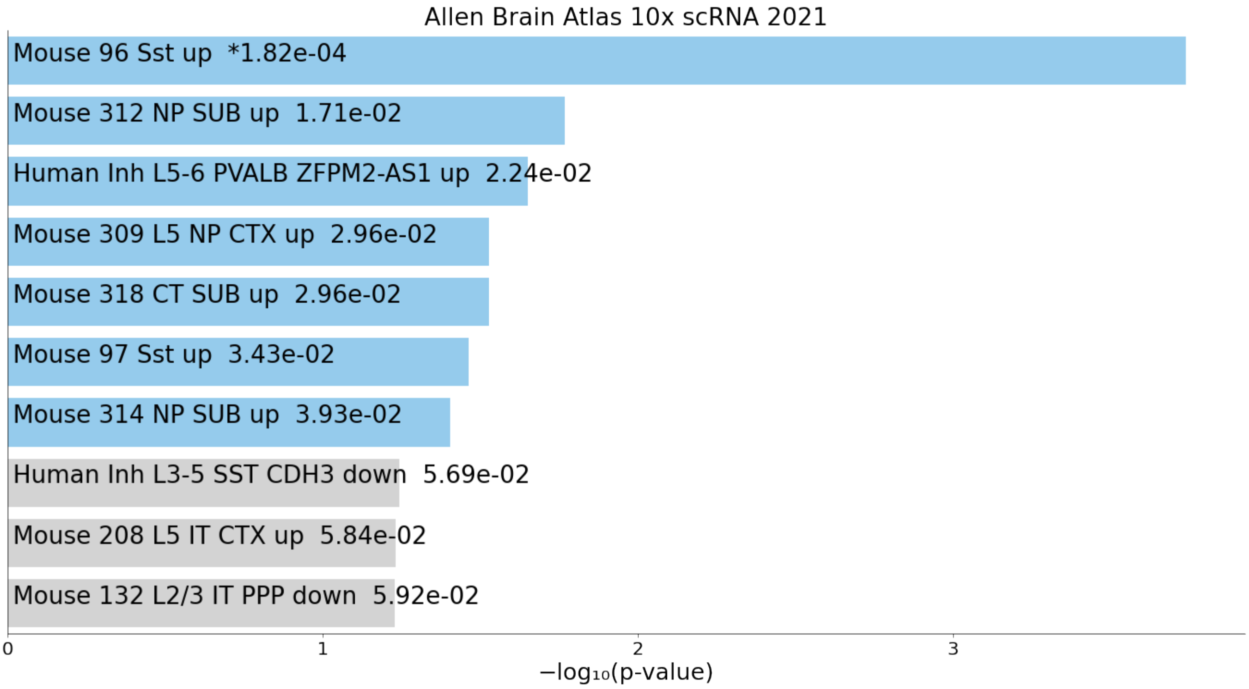


Figure S4 right_cerebral_cortex_up1

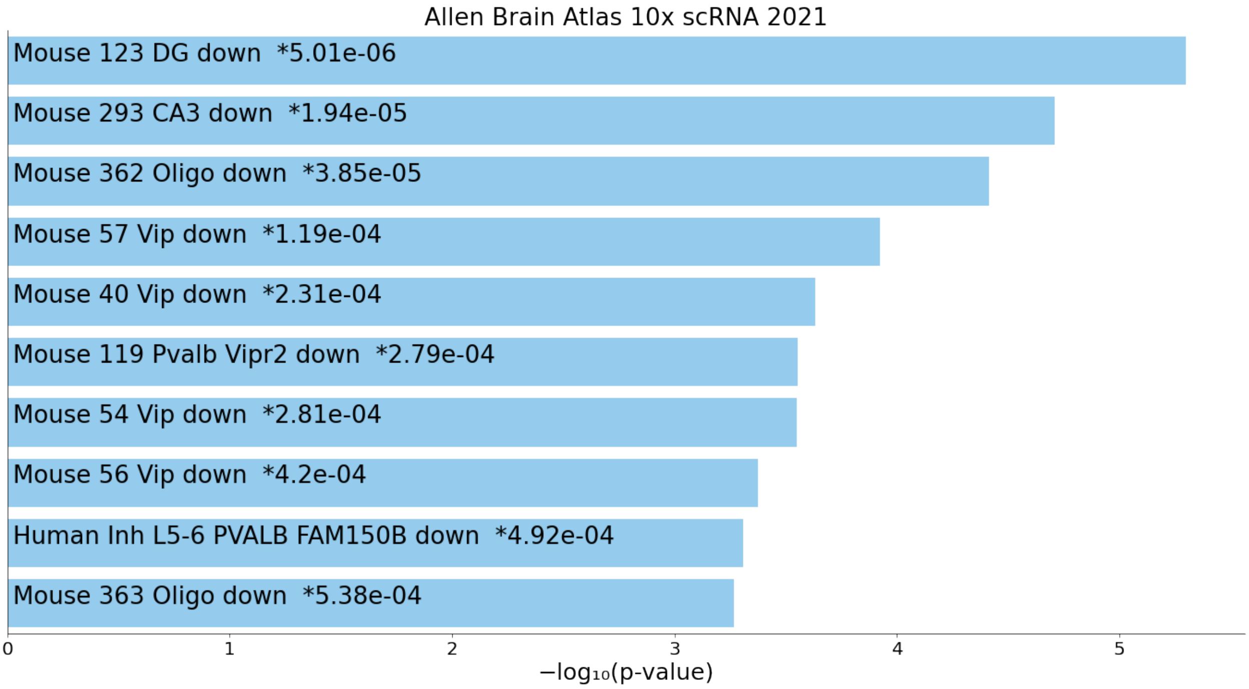


Figure S5 right_cerebral_cortex_down1(brain)

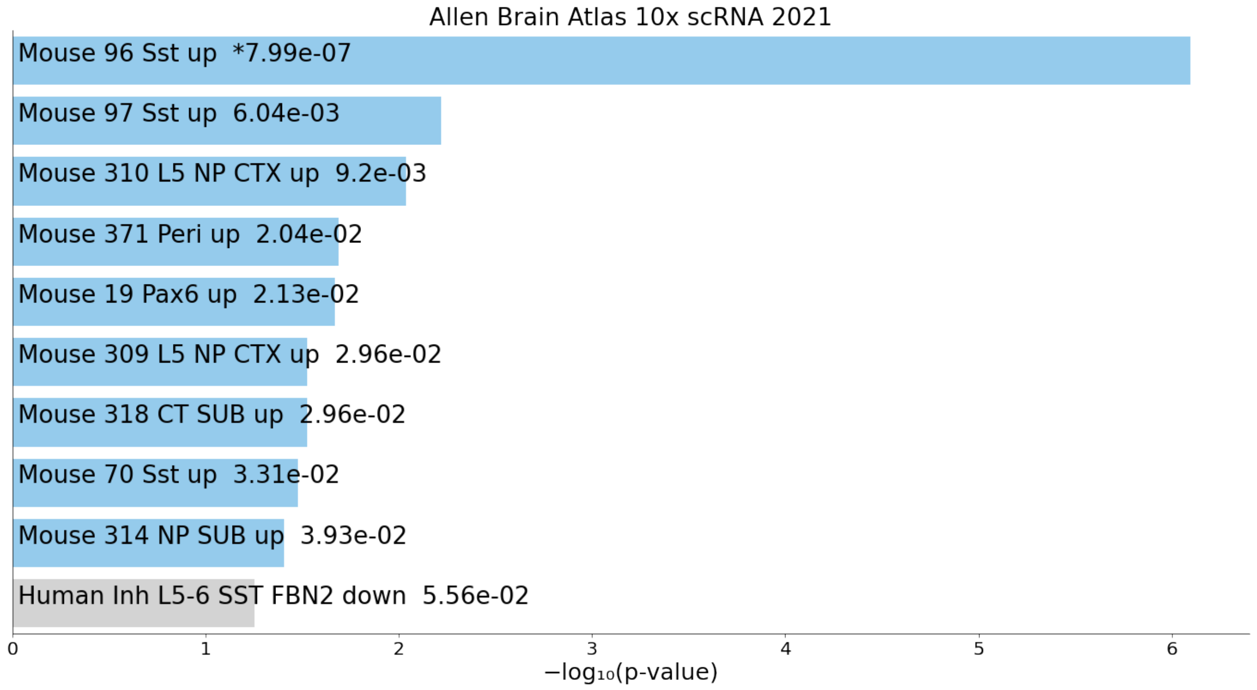


Figure S6 hippocampal_layer_up1

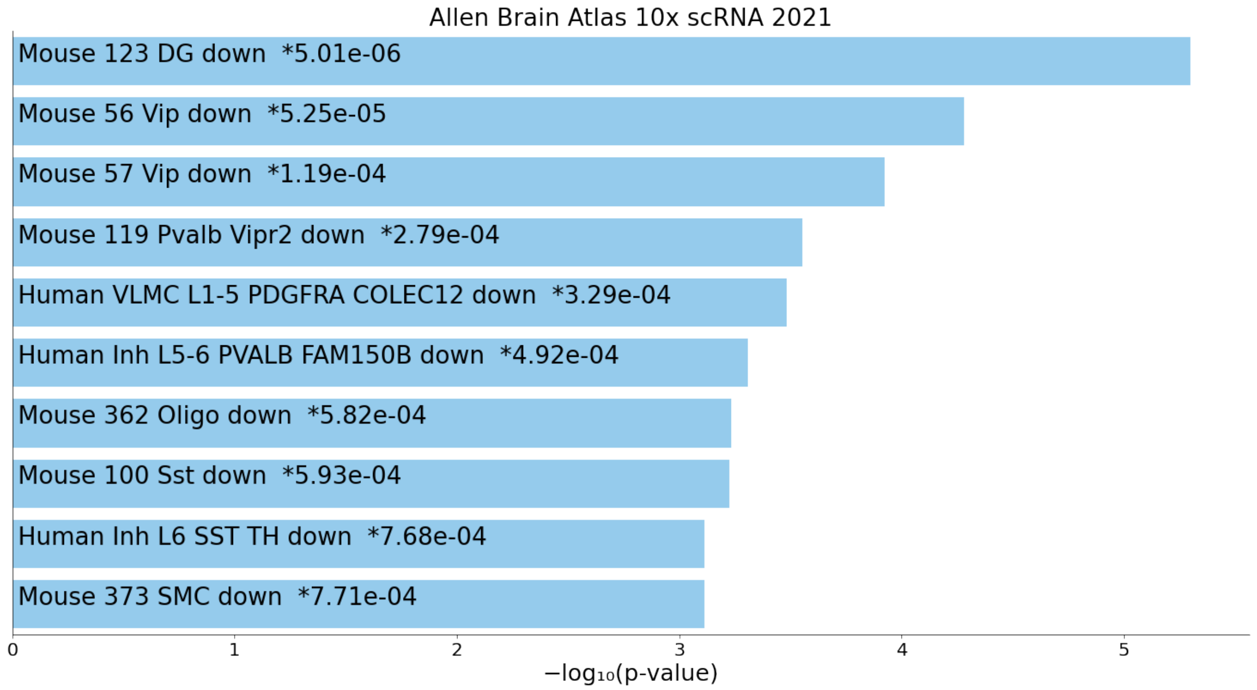


Figure S7 hippocampal_layer_down1(brain up)

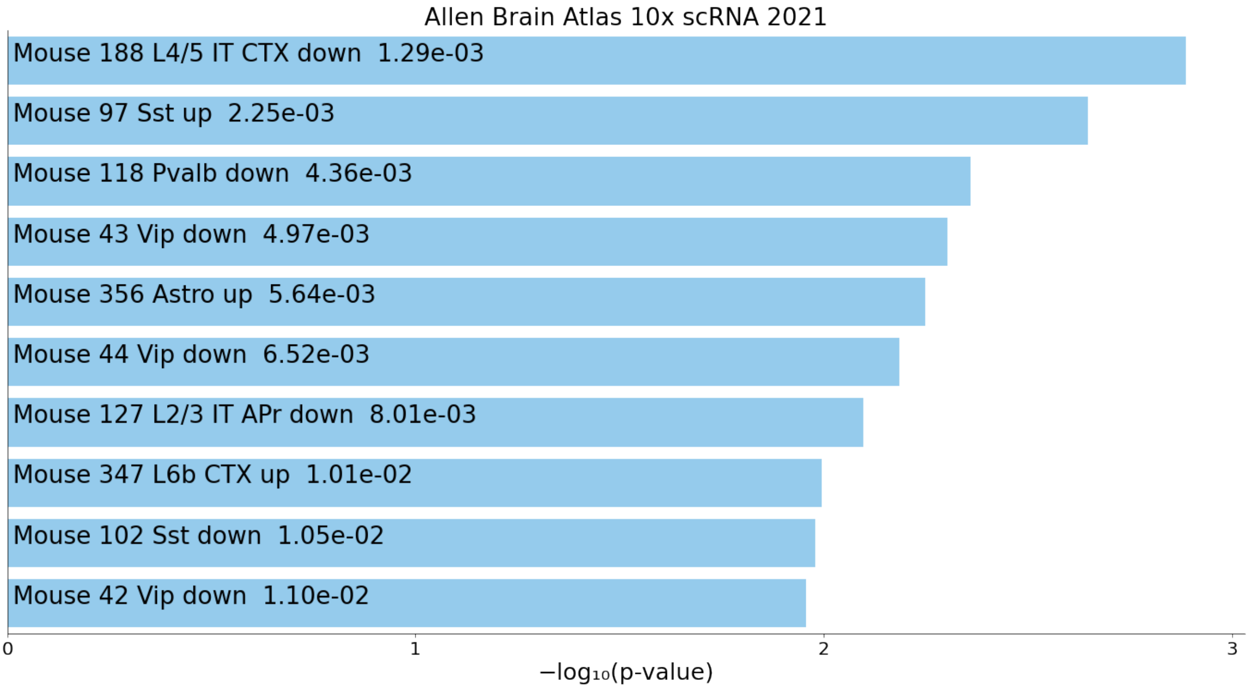


Figure S8 frontal_cortex_up1

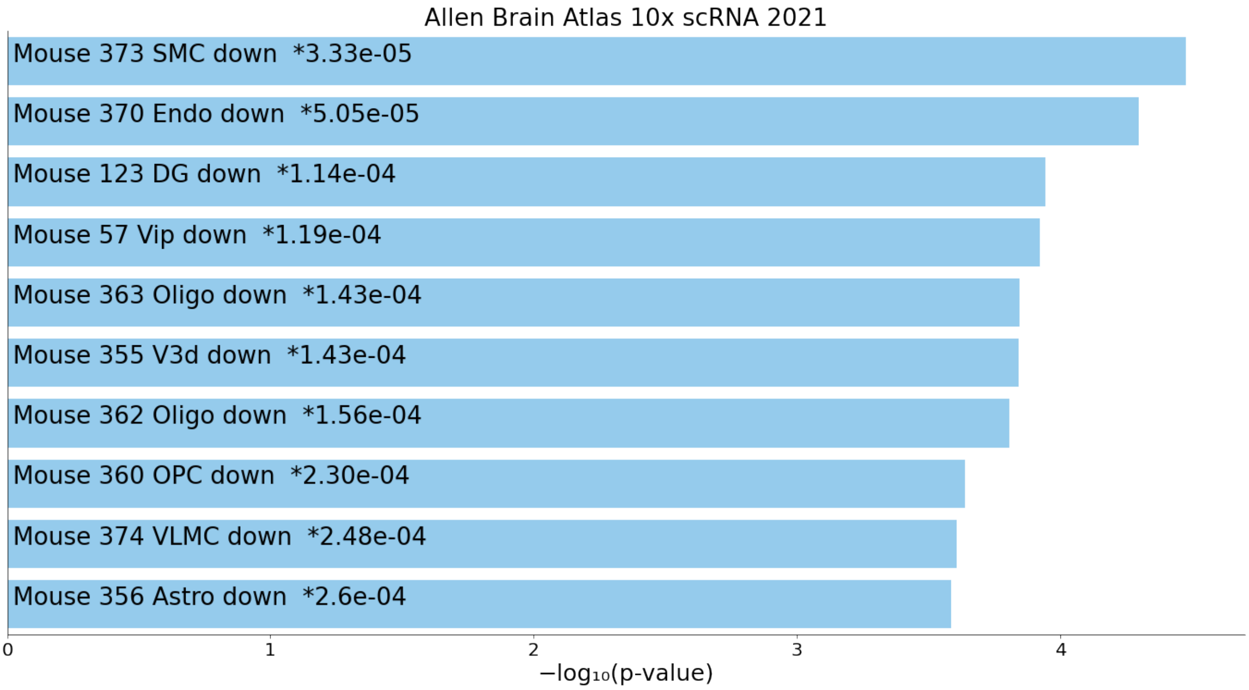


Figure S9 frontal_cortex_down1(brain up)

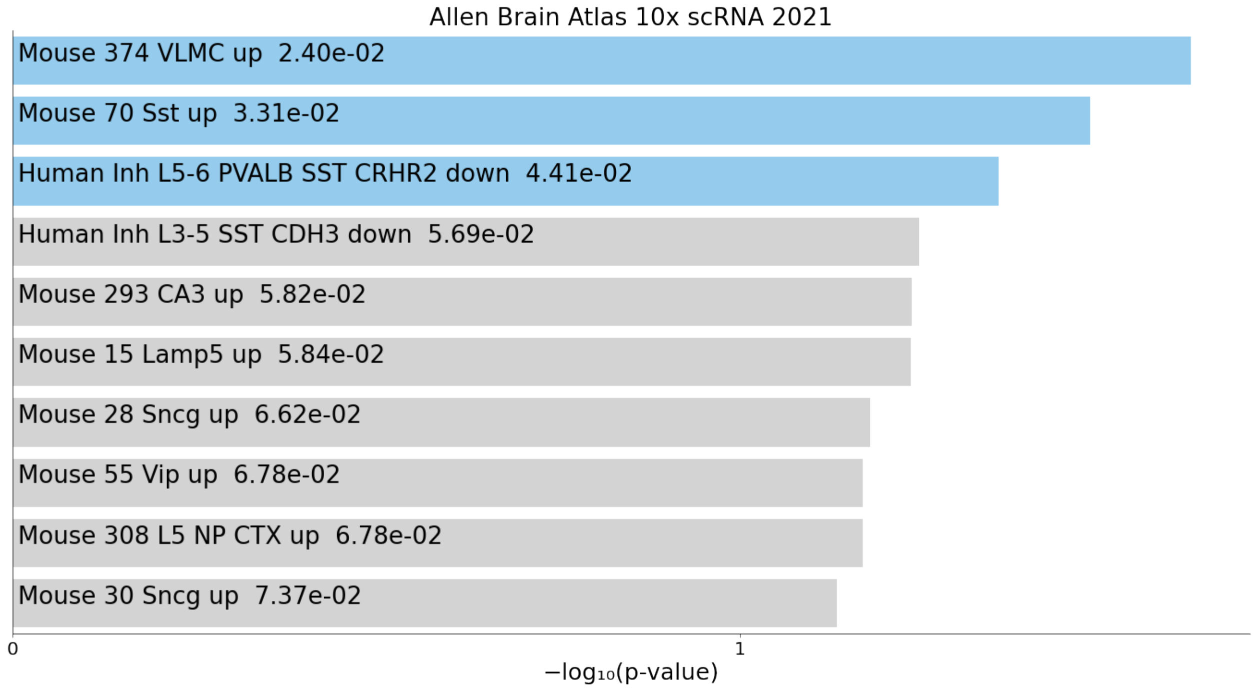


Figure S10 left_cerebral_cortex_up2

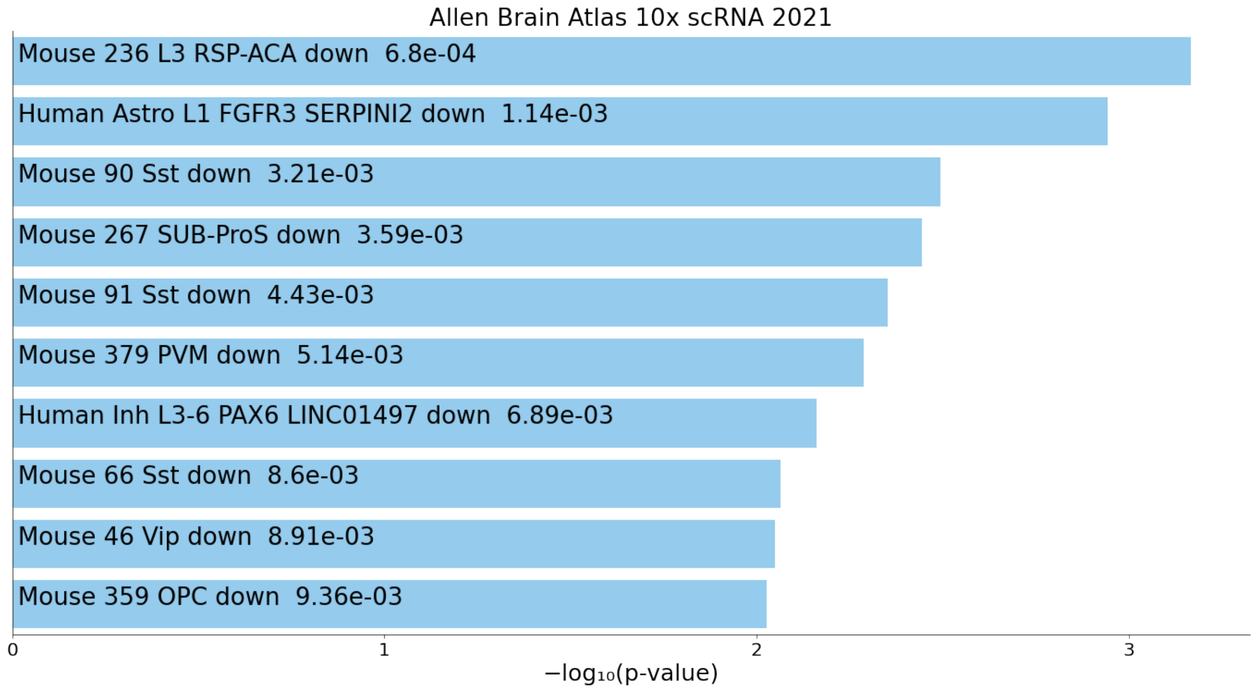


Figure S11 left_cerebral_cortex_down2(brain up)

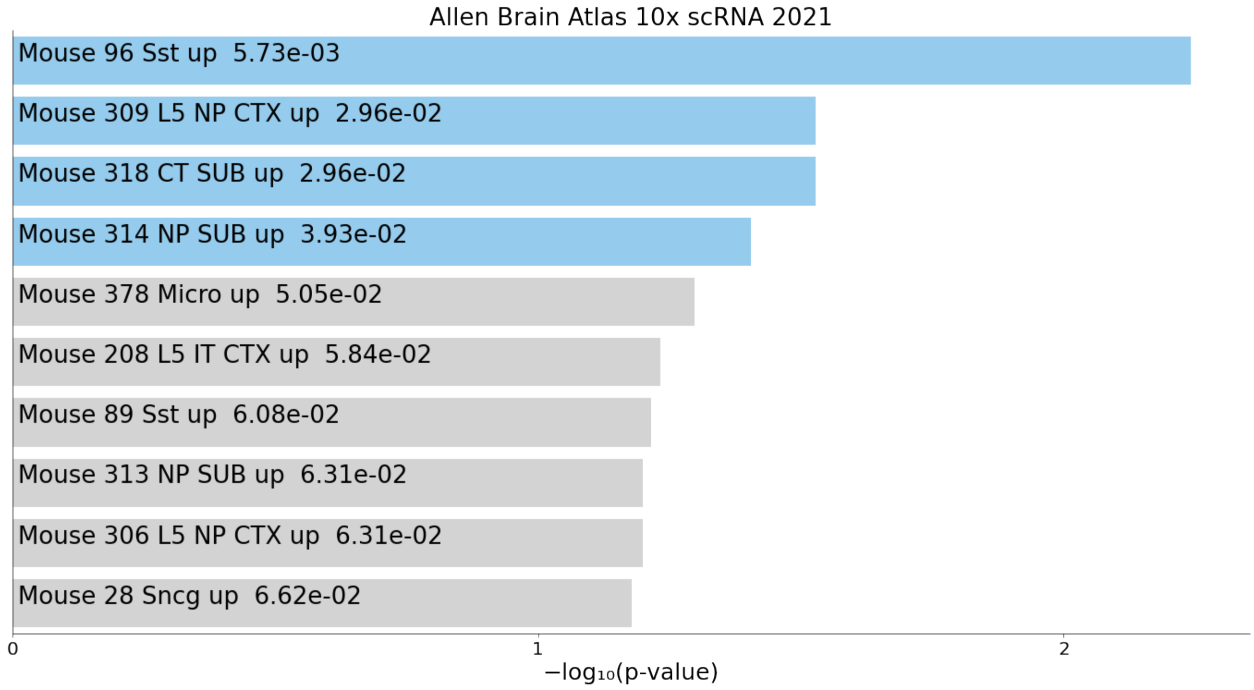


Figure S12 right_cerebral_cortex_up2

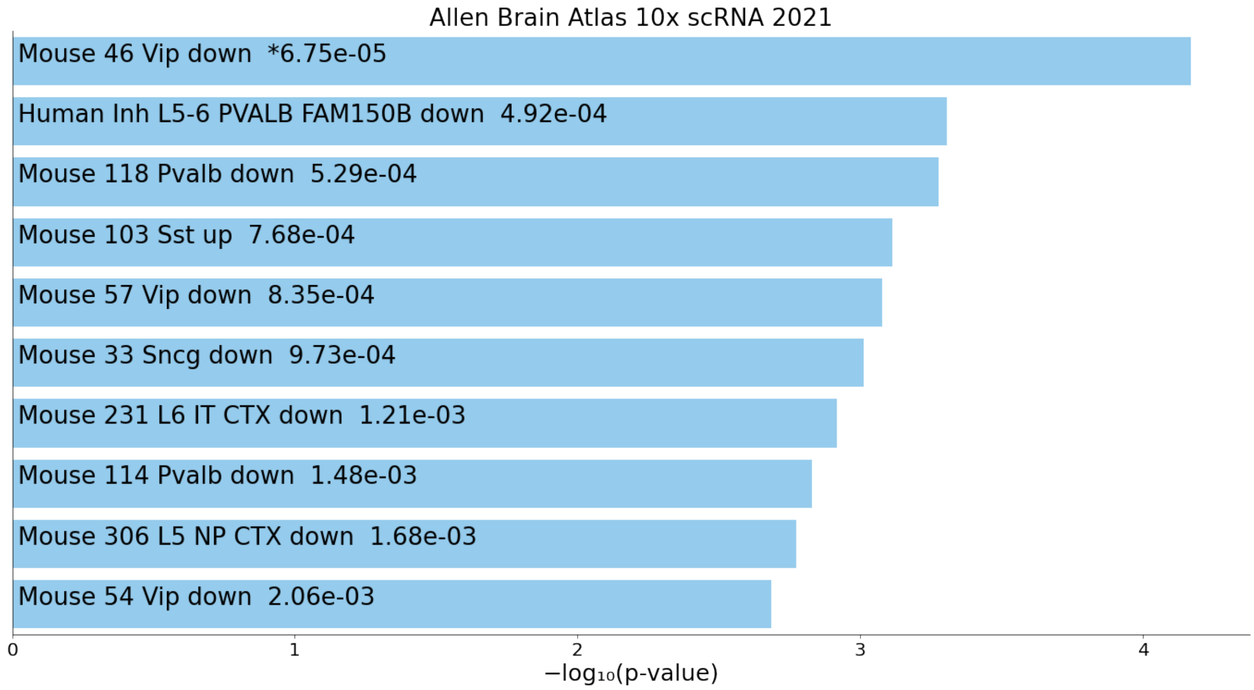


Figure S13 right_cerebral_cortex_down2(brain up)

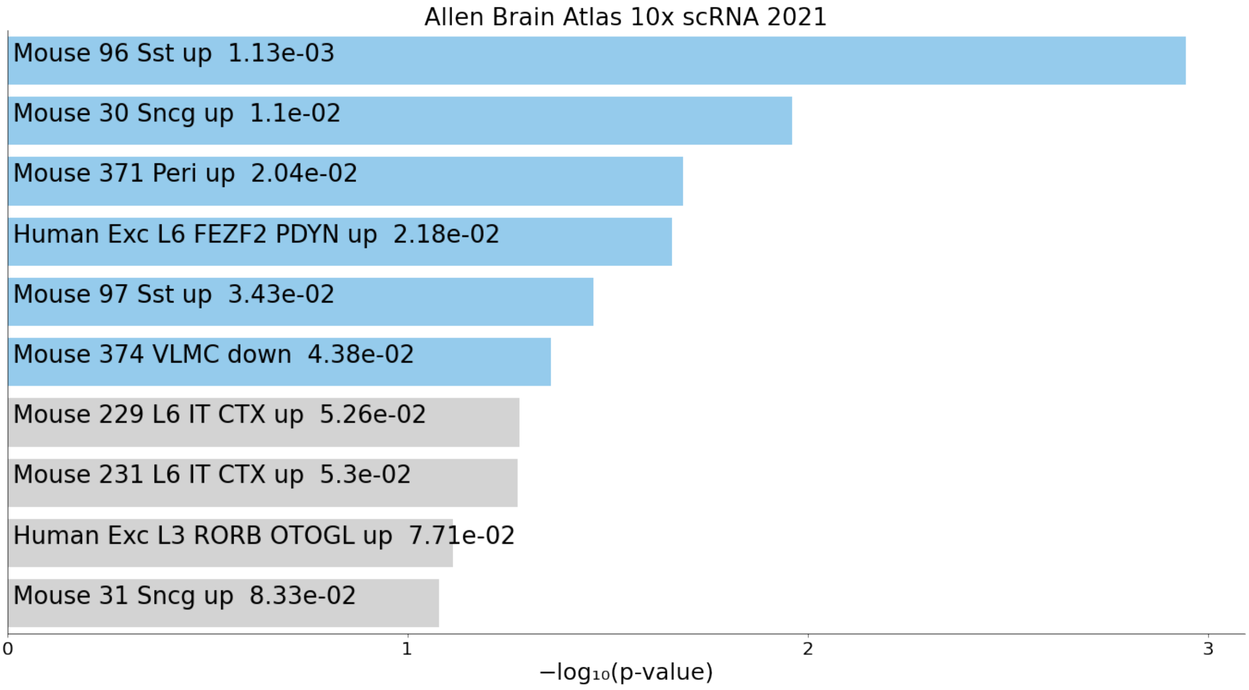


Figure S14 hippocampal_layer_up2

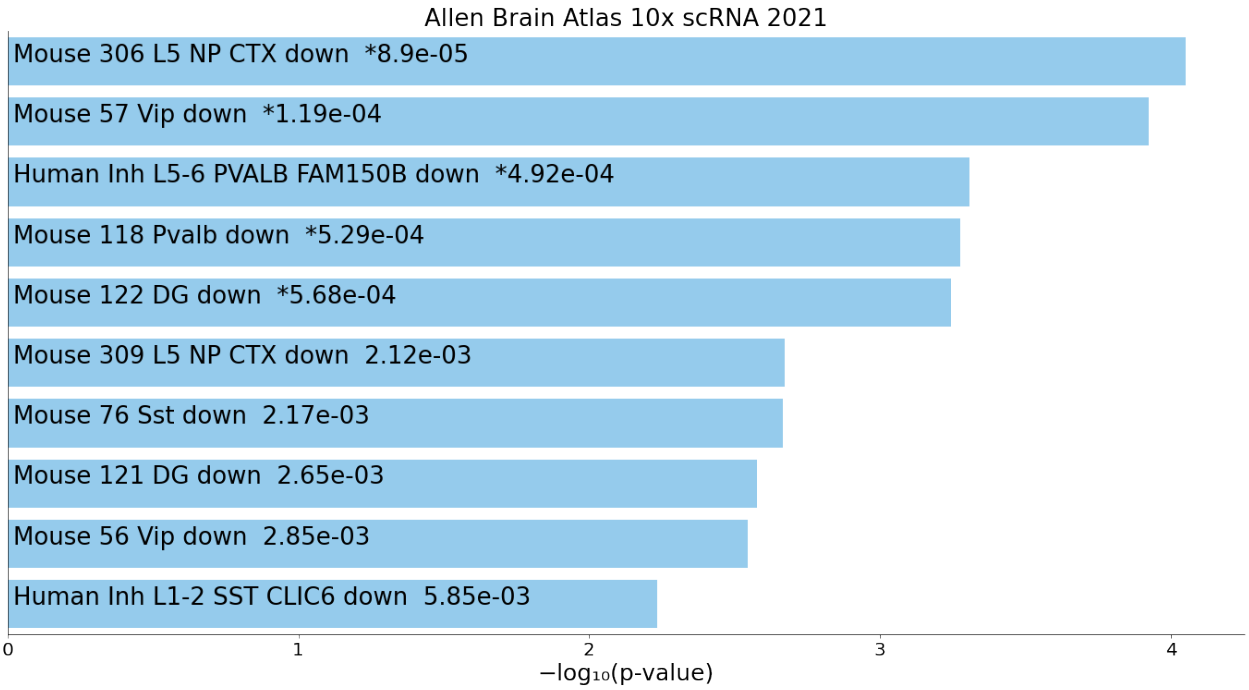


Figure S15 hippocampal_layer_down2(brain up)

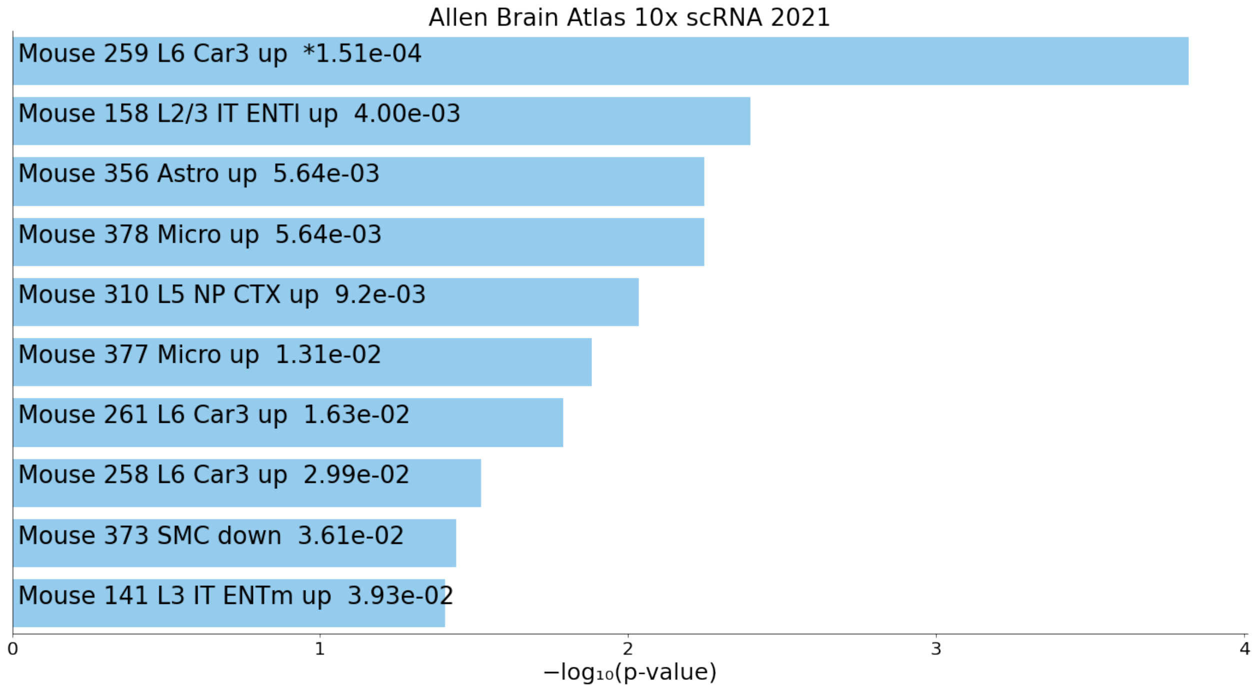


Figure S16 frontal_cortex_up2

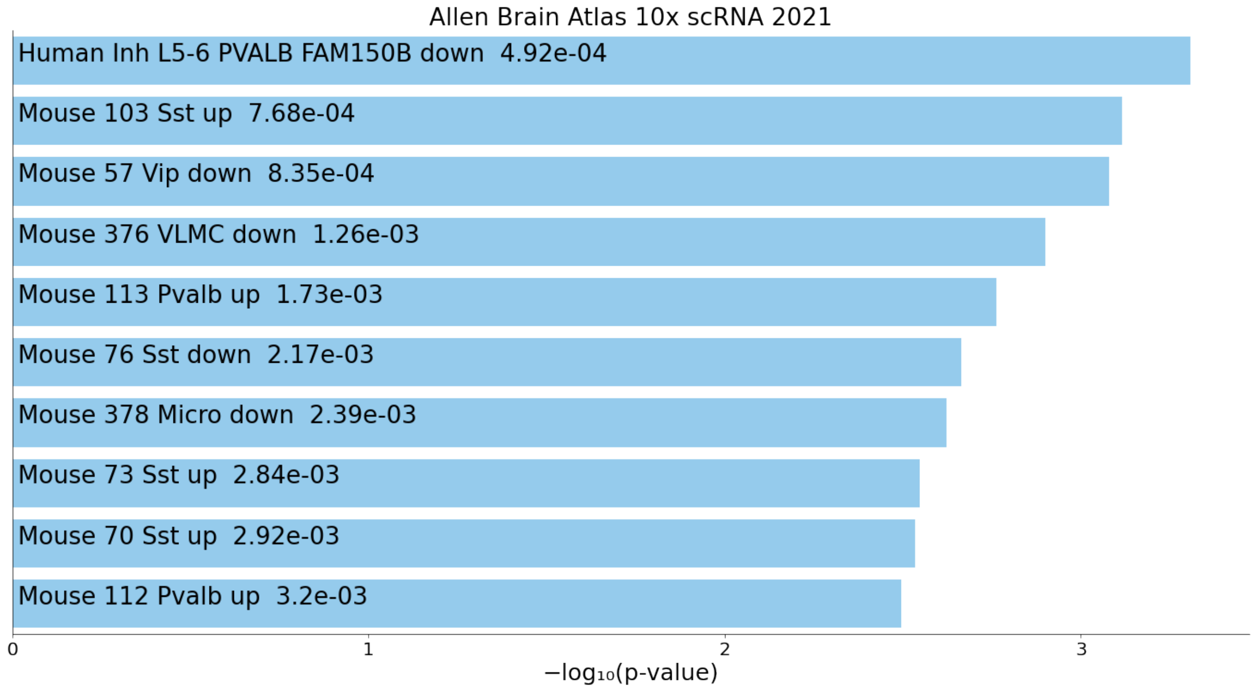


Figure S17 frontal_cortex_down2(brain up)

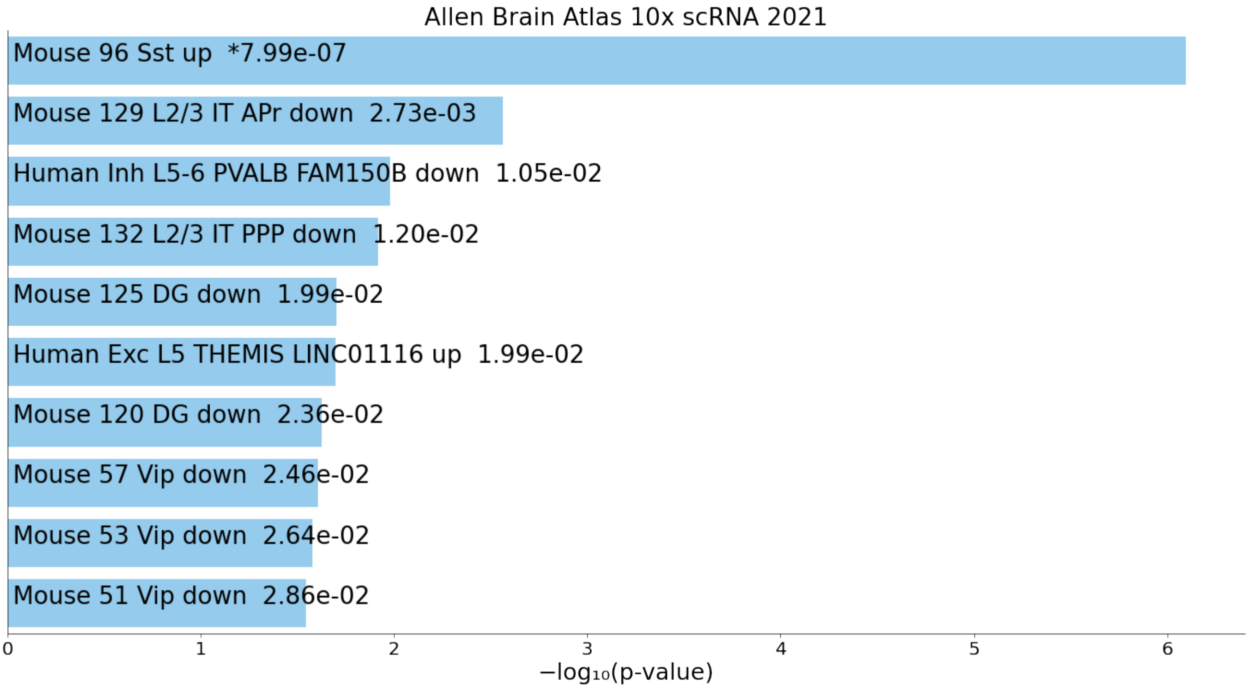


Figure S18 left_cerebral_cortex_up

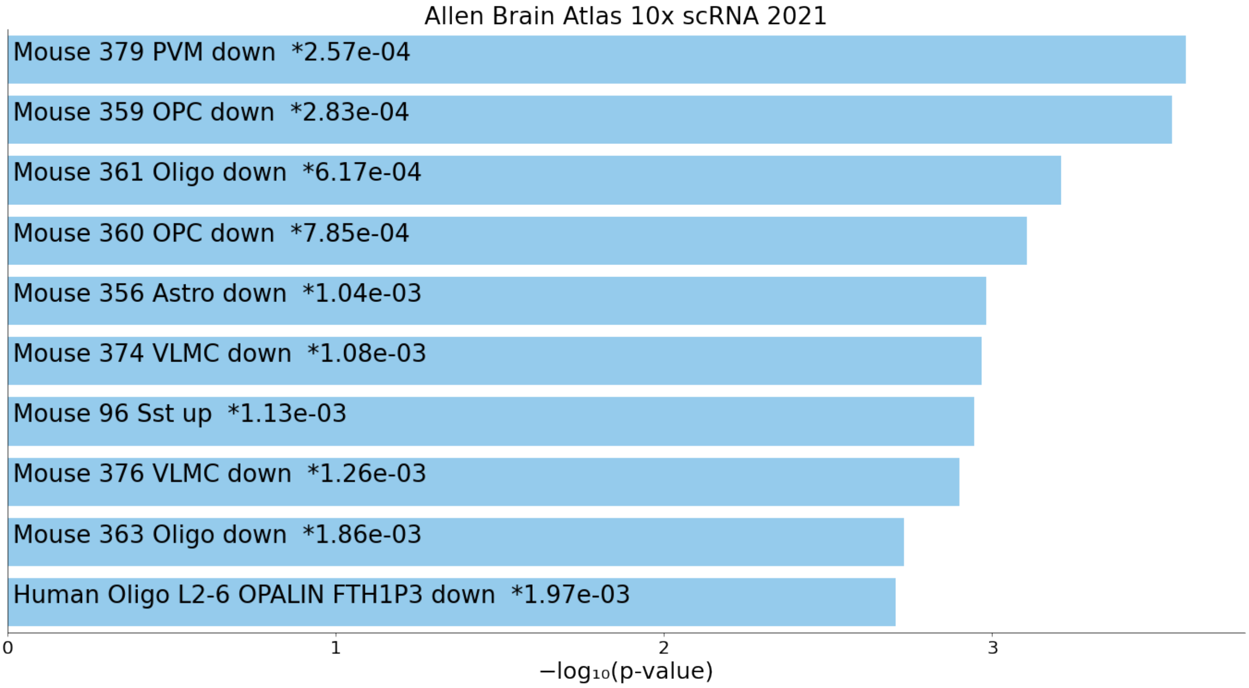


Figure S19 left_cerebral_cortex_down(brain up)

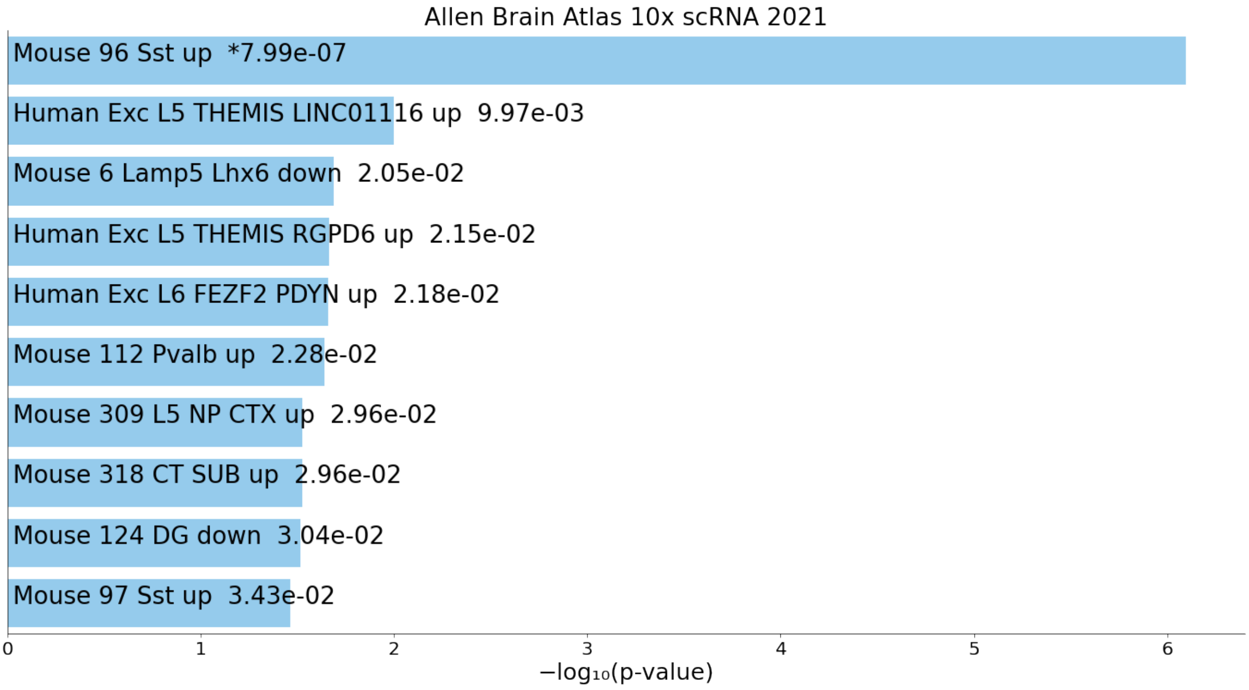


Figure S20 right_cerebral_cortex_up

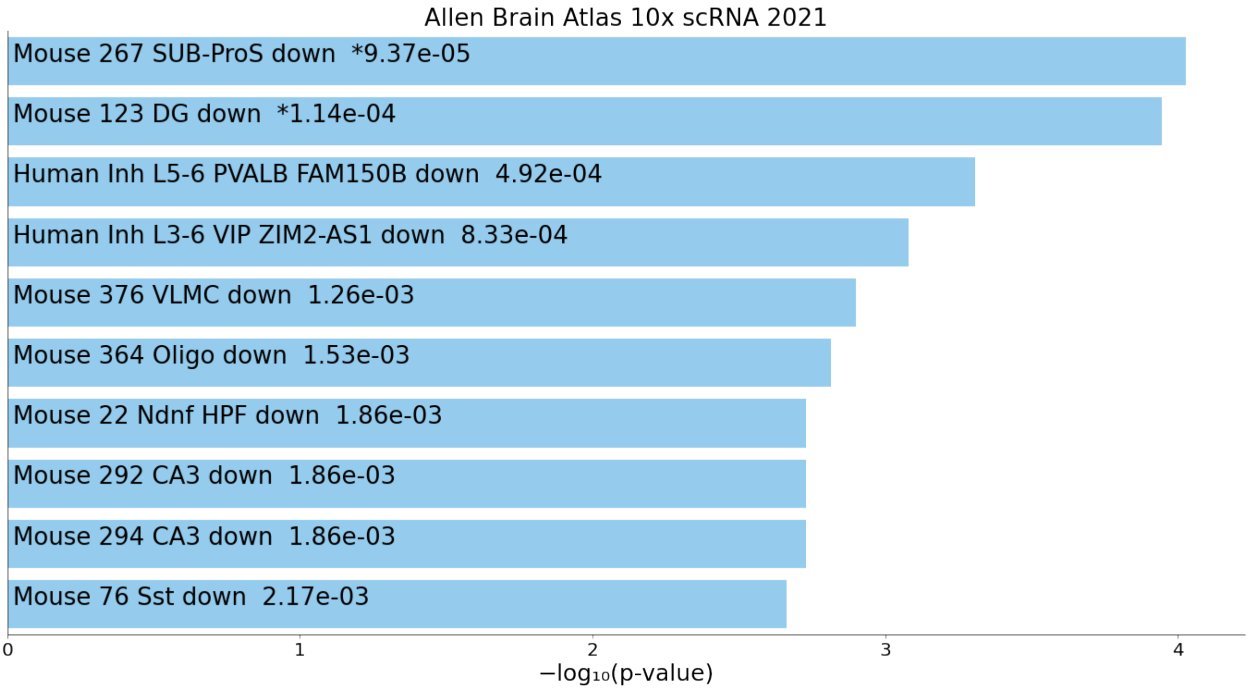


Figure S21 right_cerebral_cortex_down(brain up)

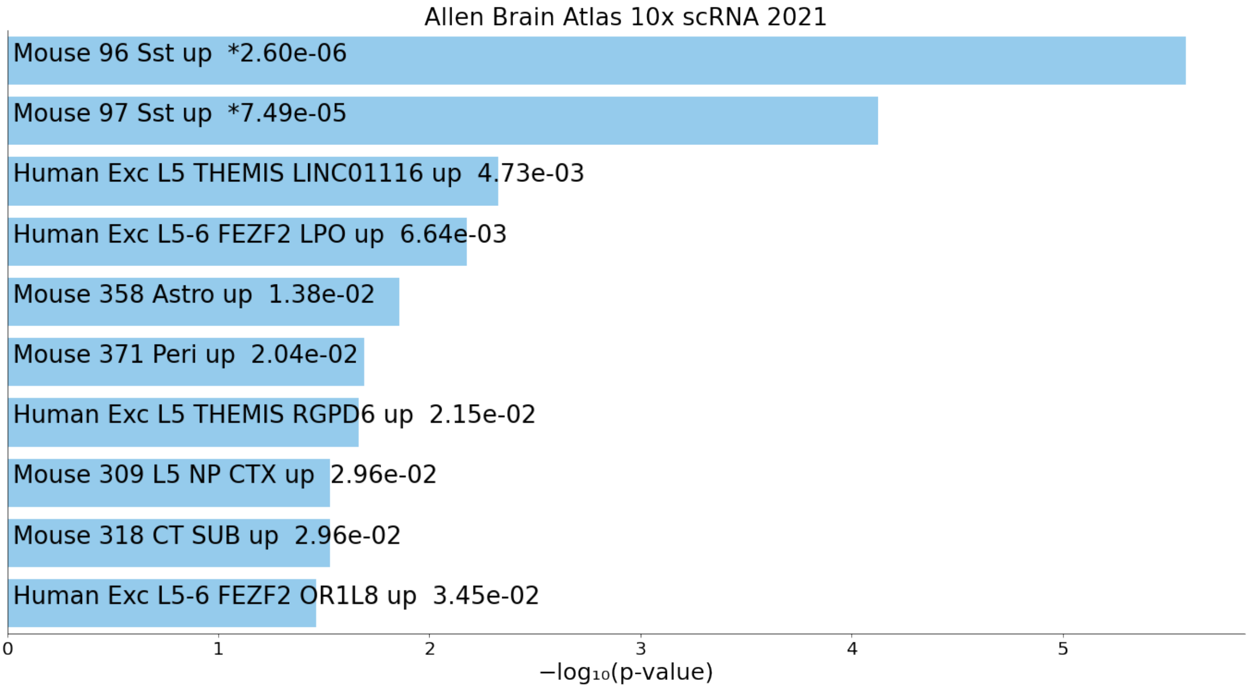


Figure S22 hippocampal_layer_up

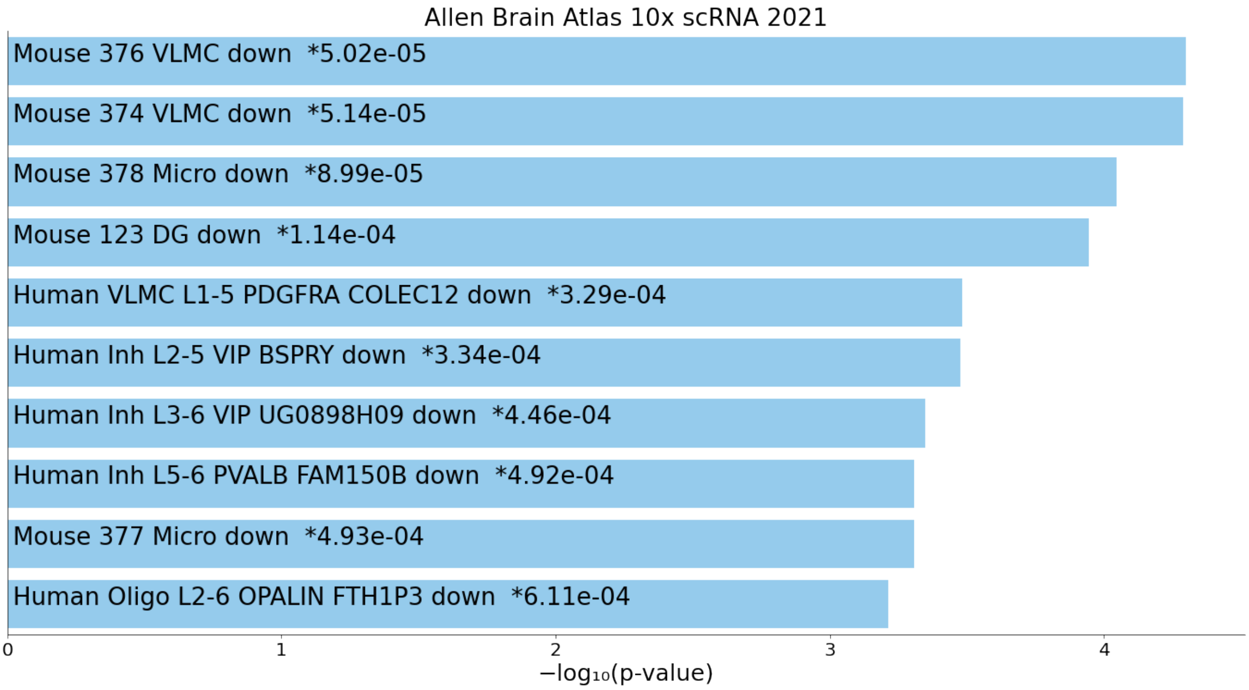


Figure S23 hippocampal_layer_down(brain up)

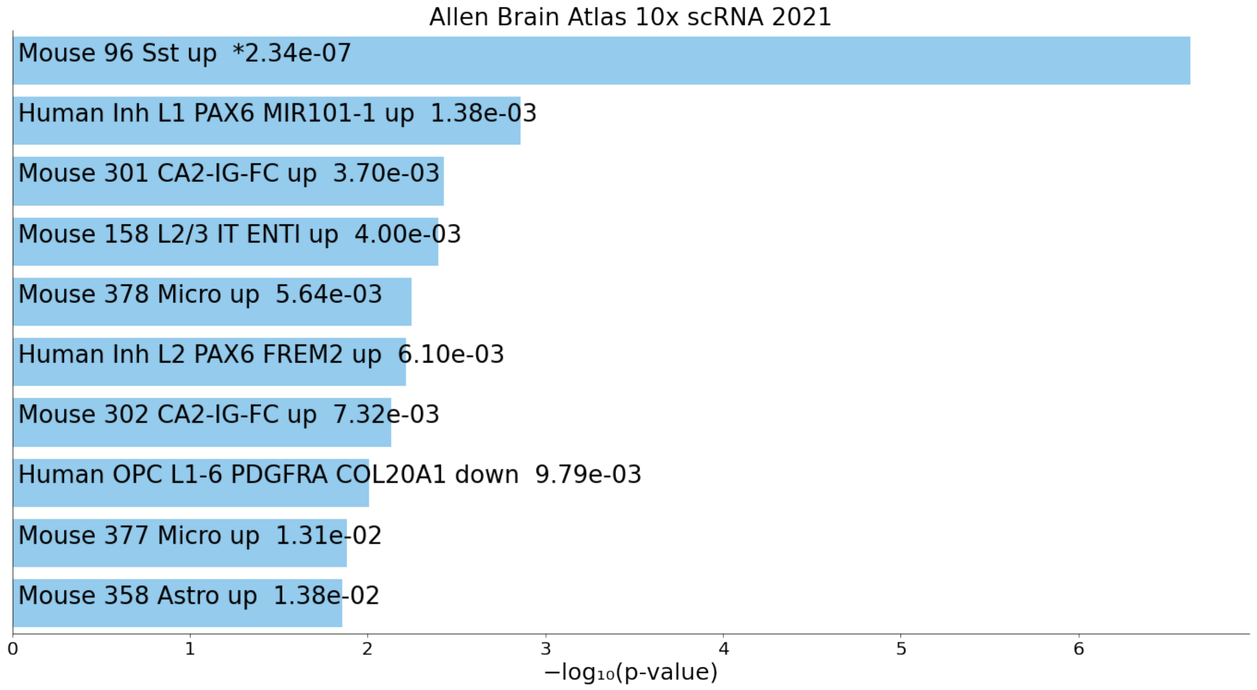


Figure S24 frontal_cortex_up

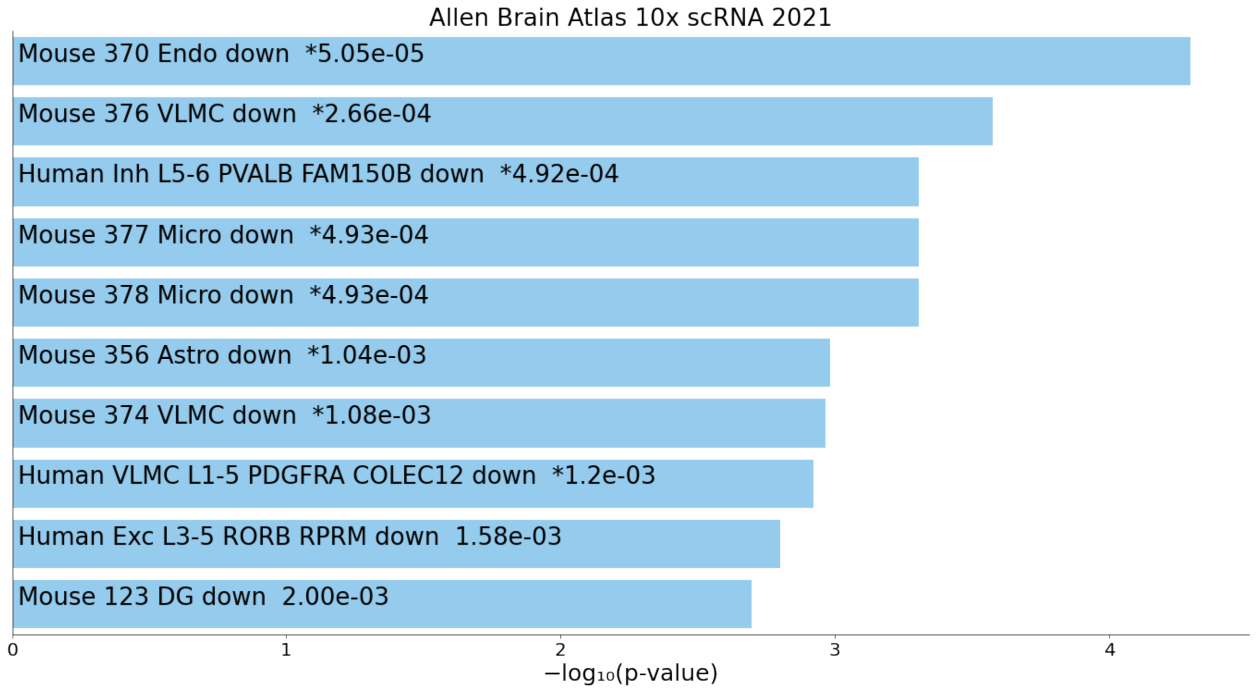


Figure S25 frontal_cortex_down(brain up)

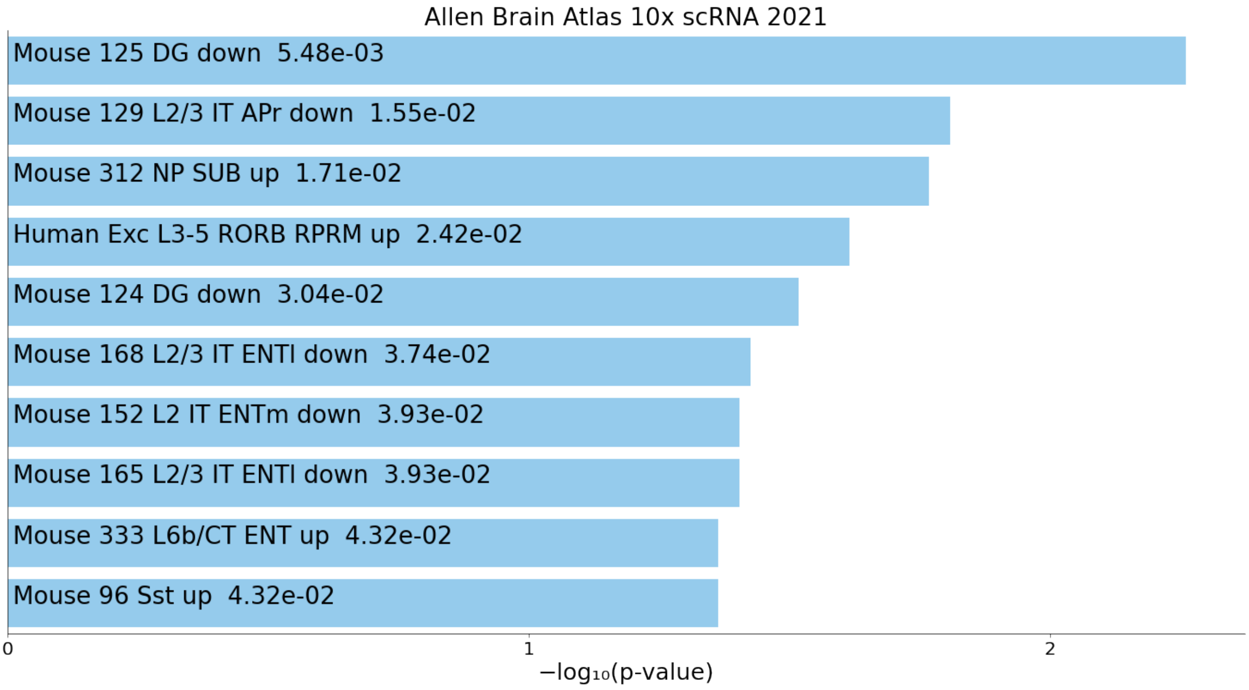


Figure S26 left_cerebral_cortex_up1

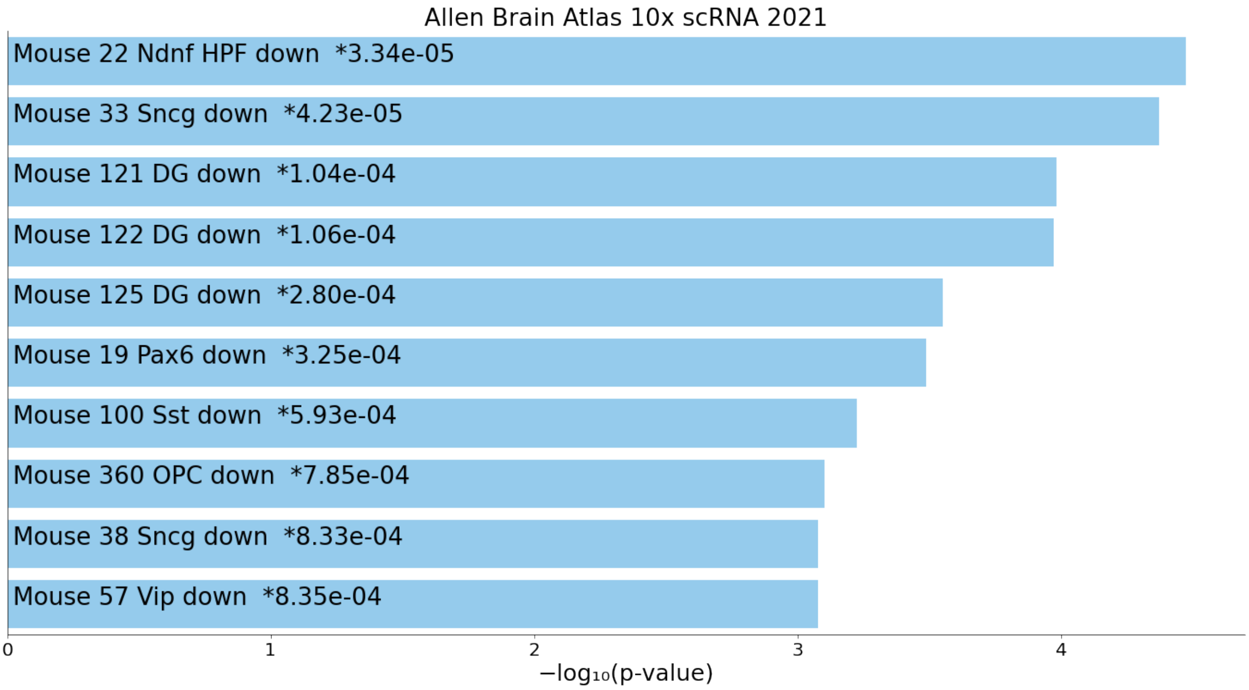


Figure S27 left_cerebral_cortex_down1(brain up)

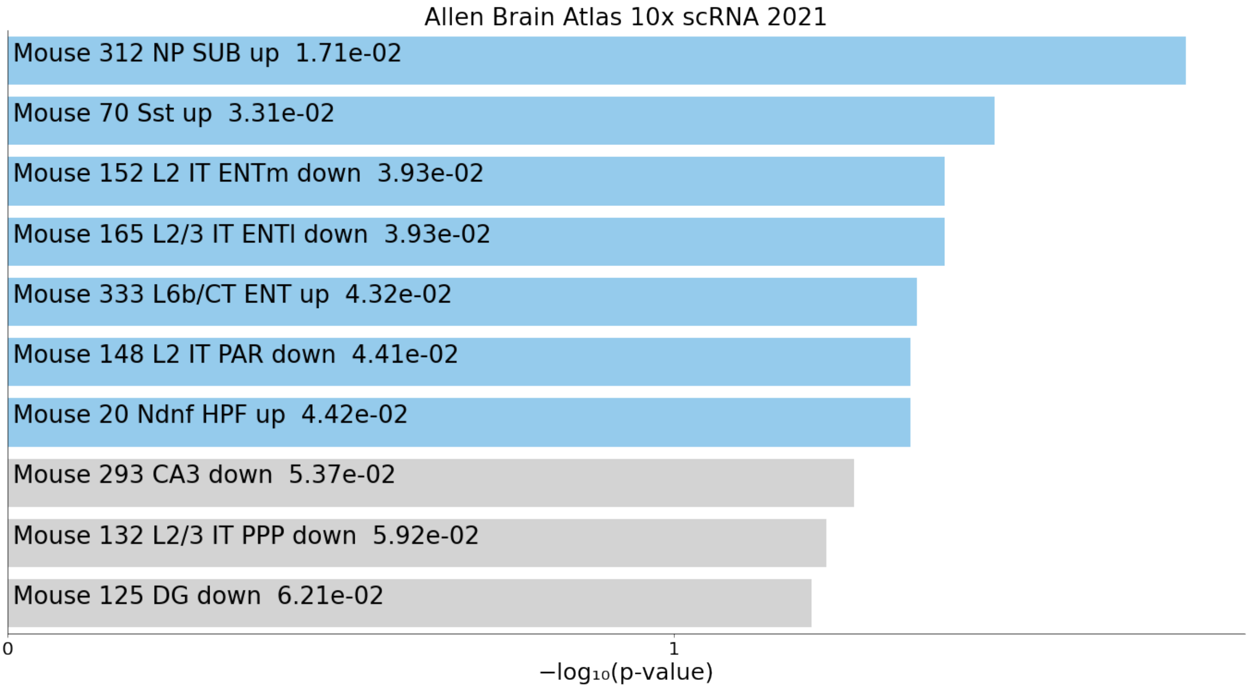


Figure S28 right_cerebral_cortex_up1

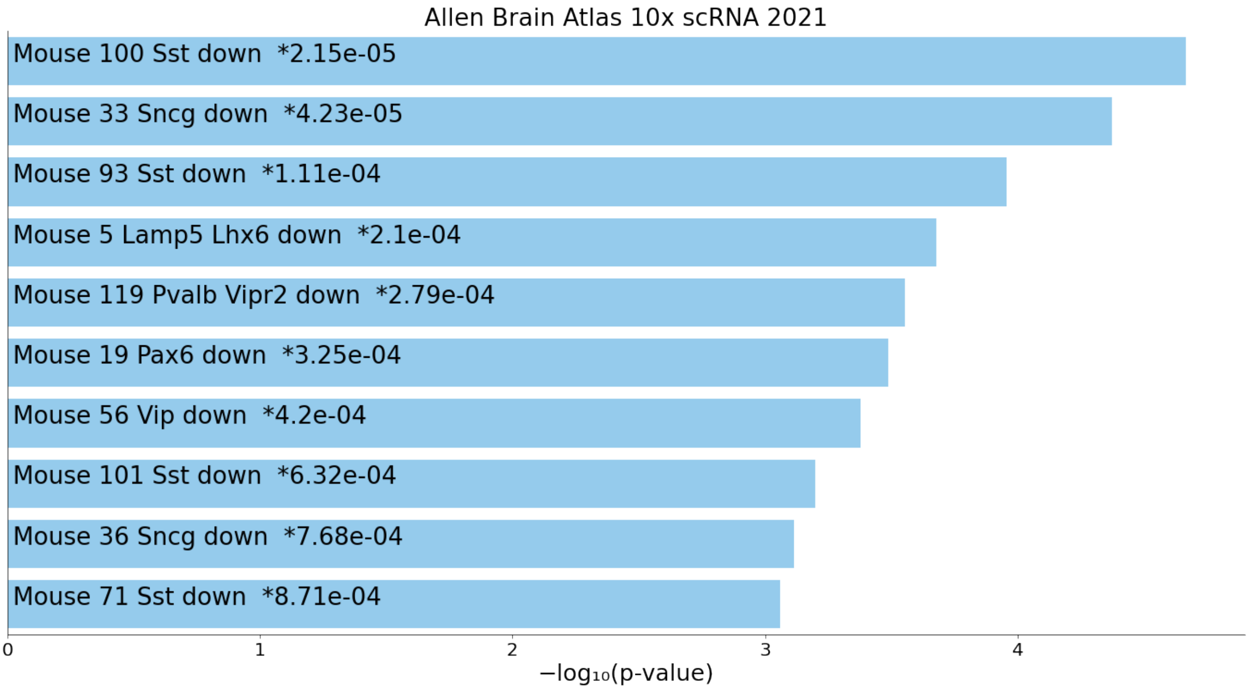


Figure S29 right_cerebral_cortex_down1(brain up)

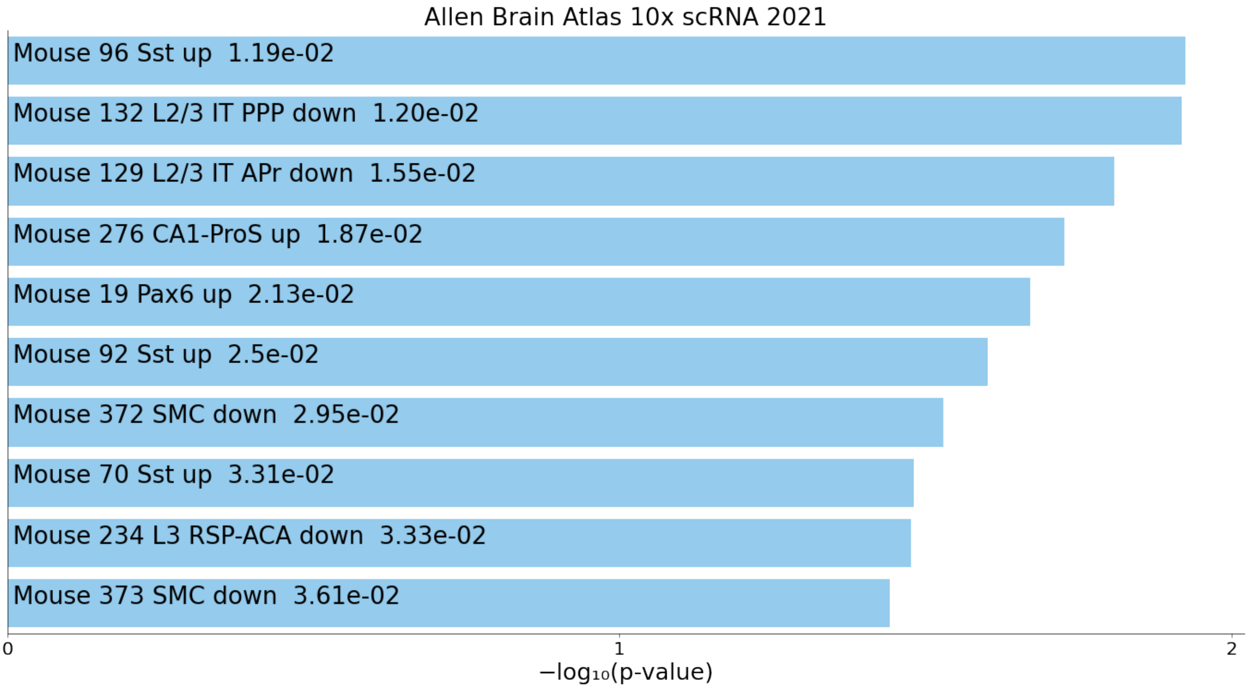


Figure S30 hippocampal_layer_up1

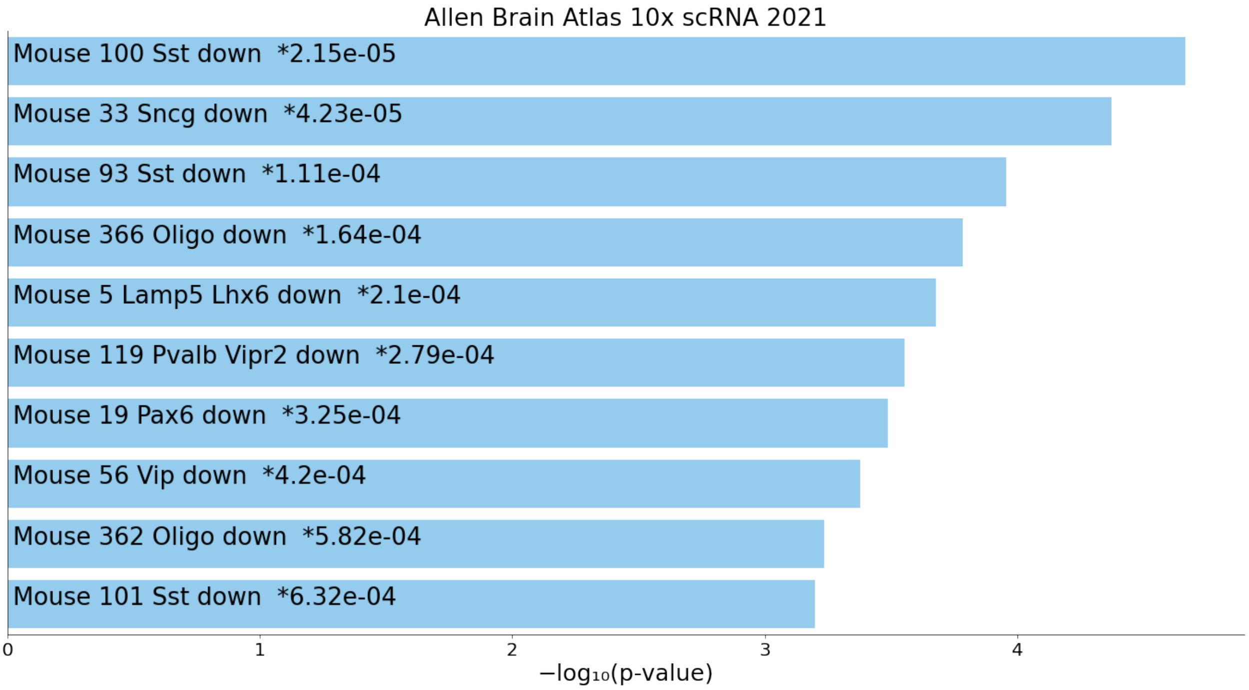


Figure S31 hippocampal_layer_down1(brain up)

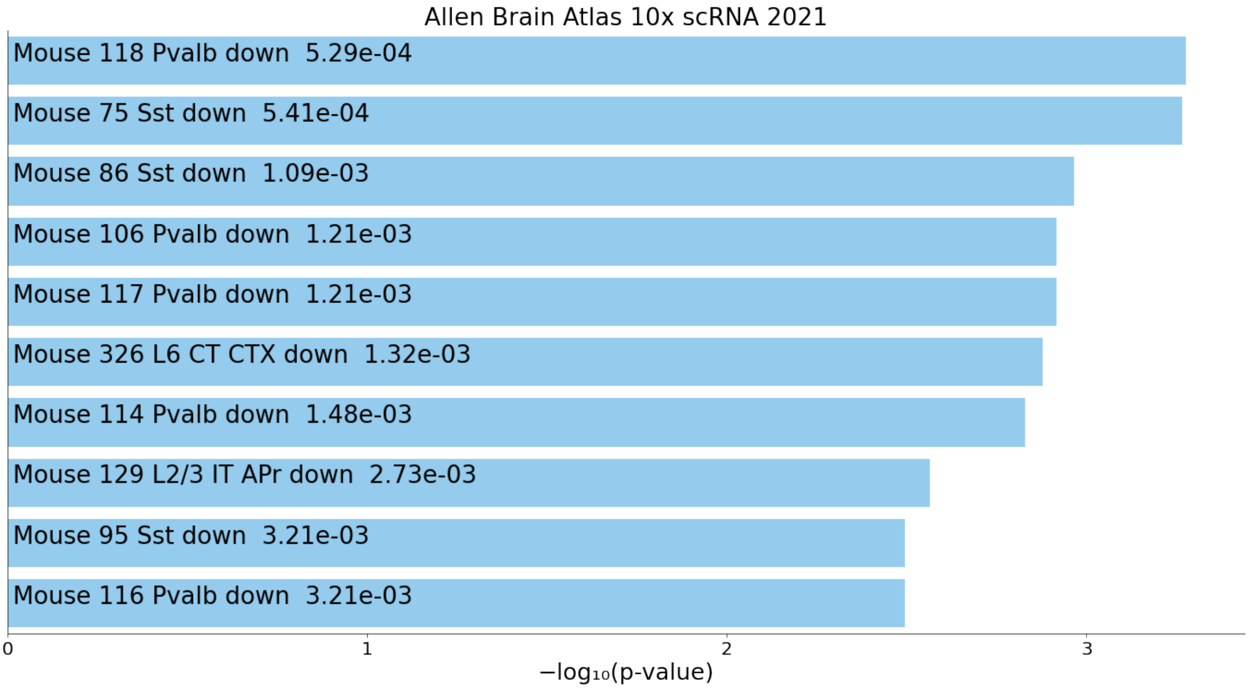


Figure S32 frontal_cortex_up1

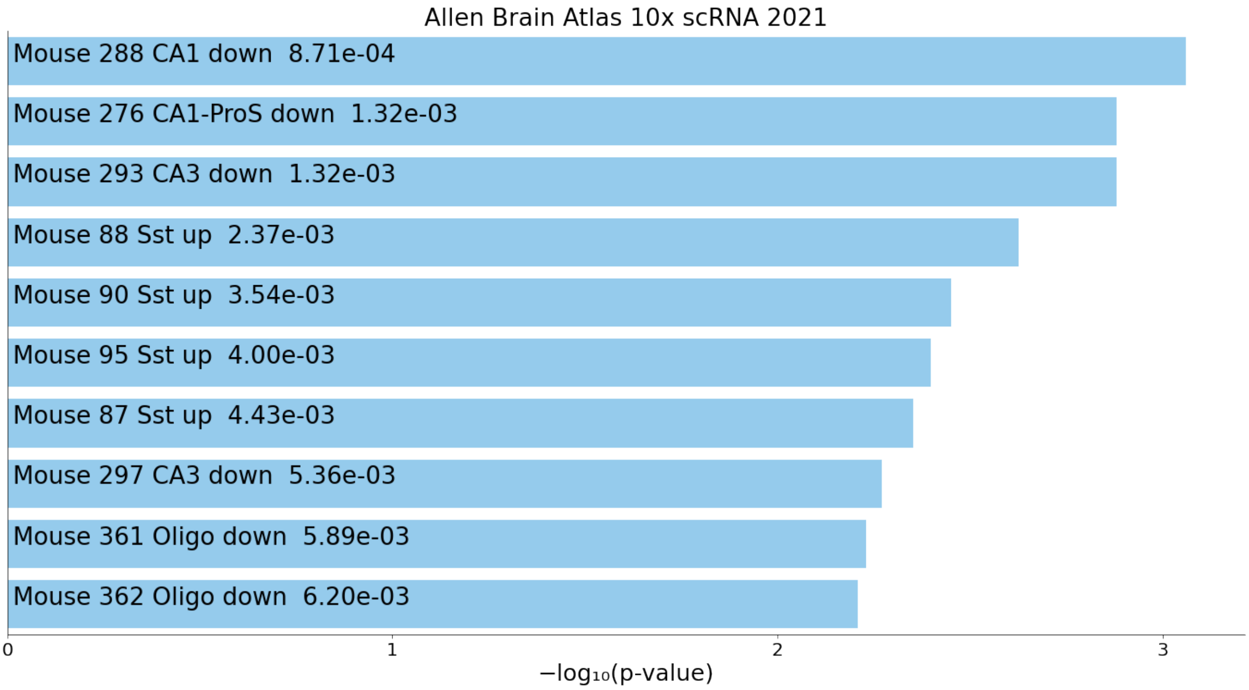


Figure S33 frontal_cortex_down1(brain up)

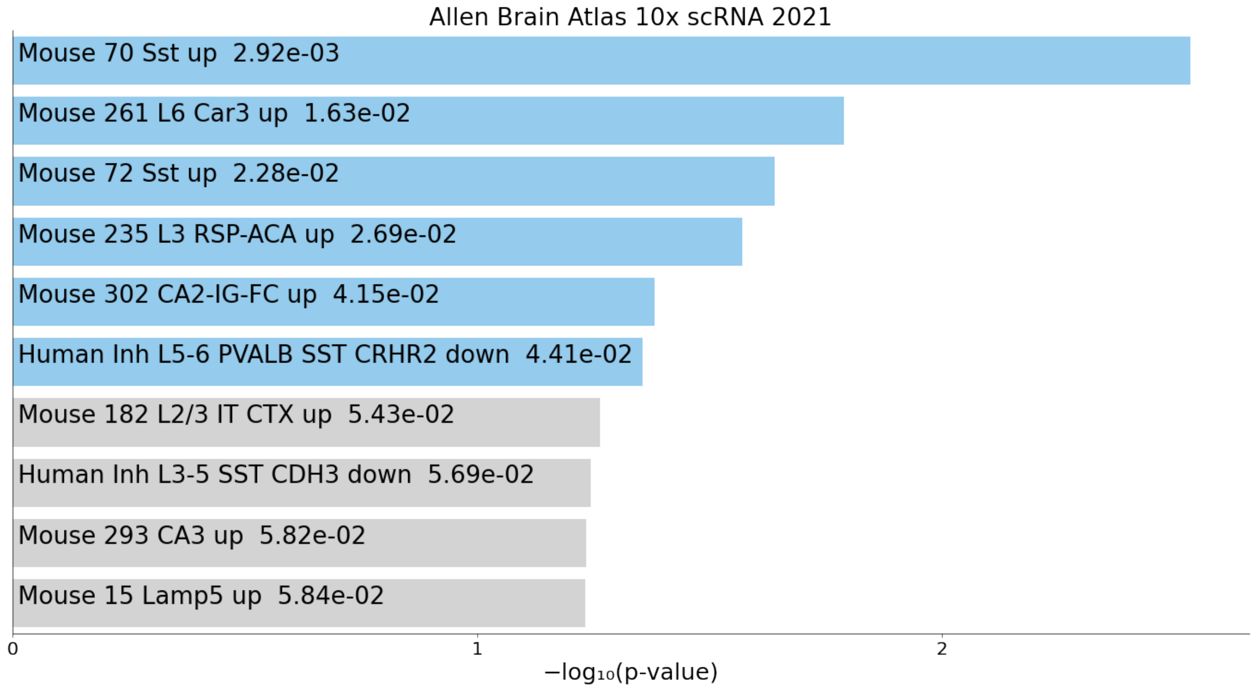


Figure S34 left_cerebral_cortex_up2

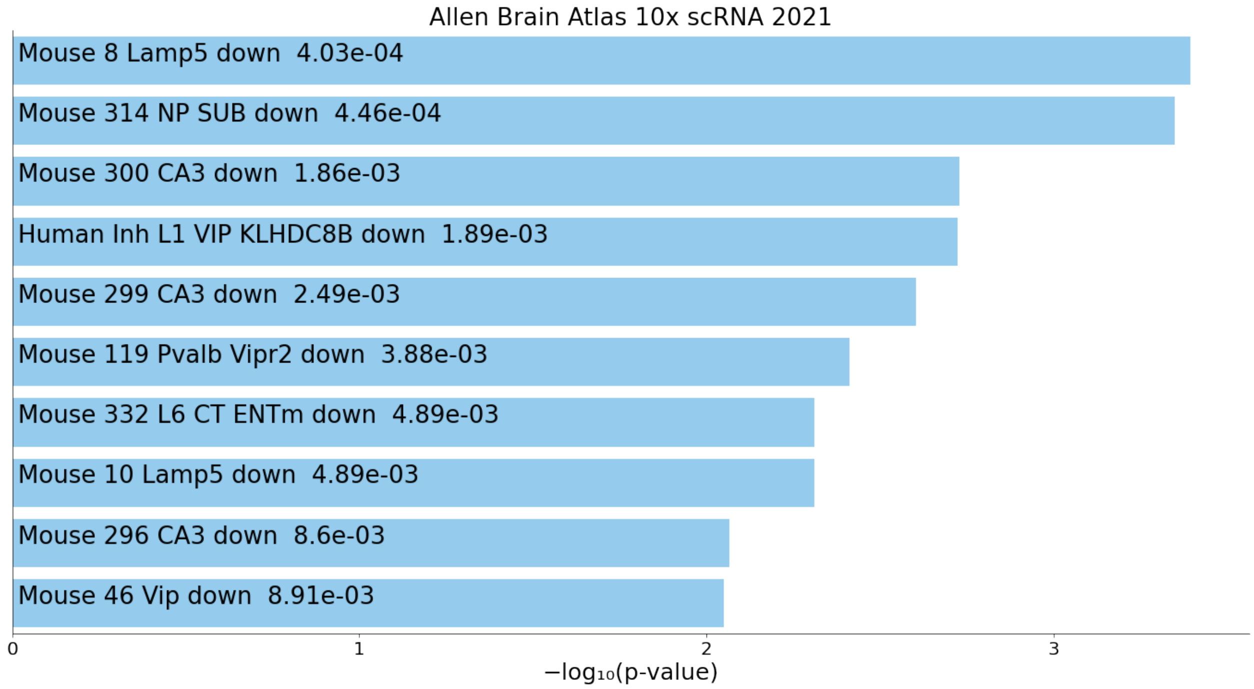


Figure S35 left_cerebral_cortex_down2(brain up)

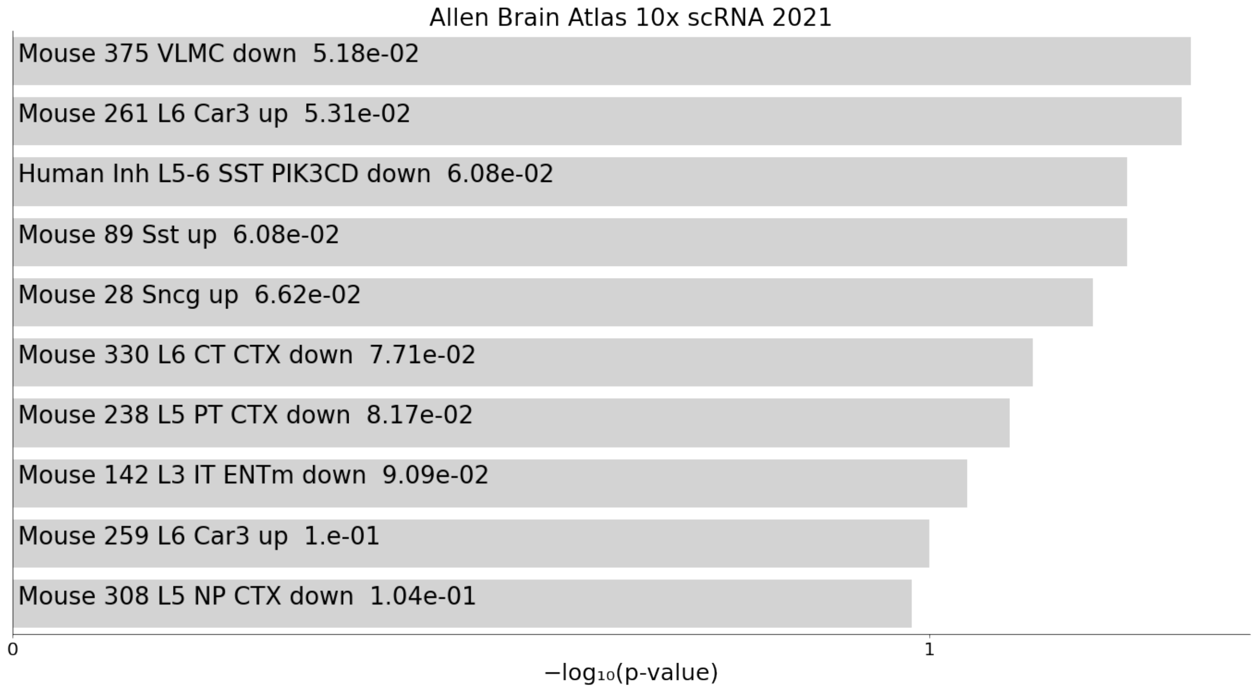


Figure S36 right_cerebral_cortex_up2

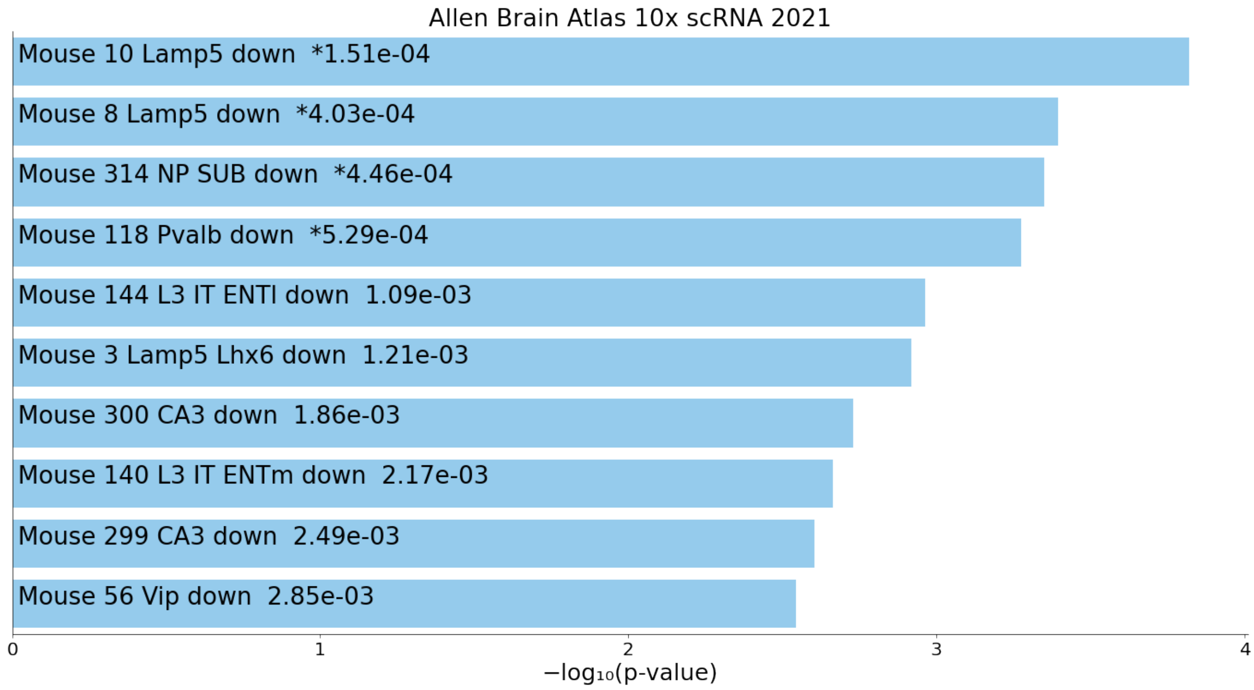


Figure S37 right_cerebral_cortex_down2(brain up)

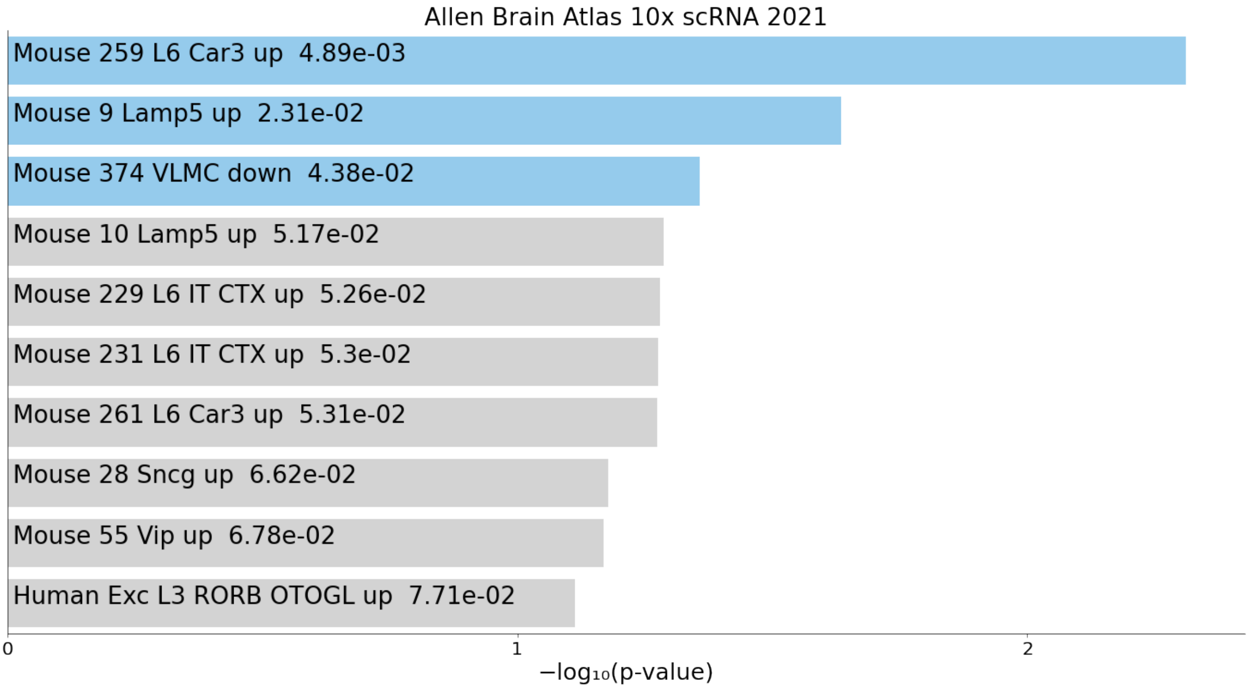


Figure S38 hippocampal_layer_up2

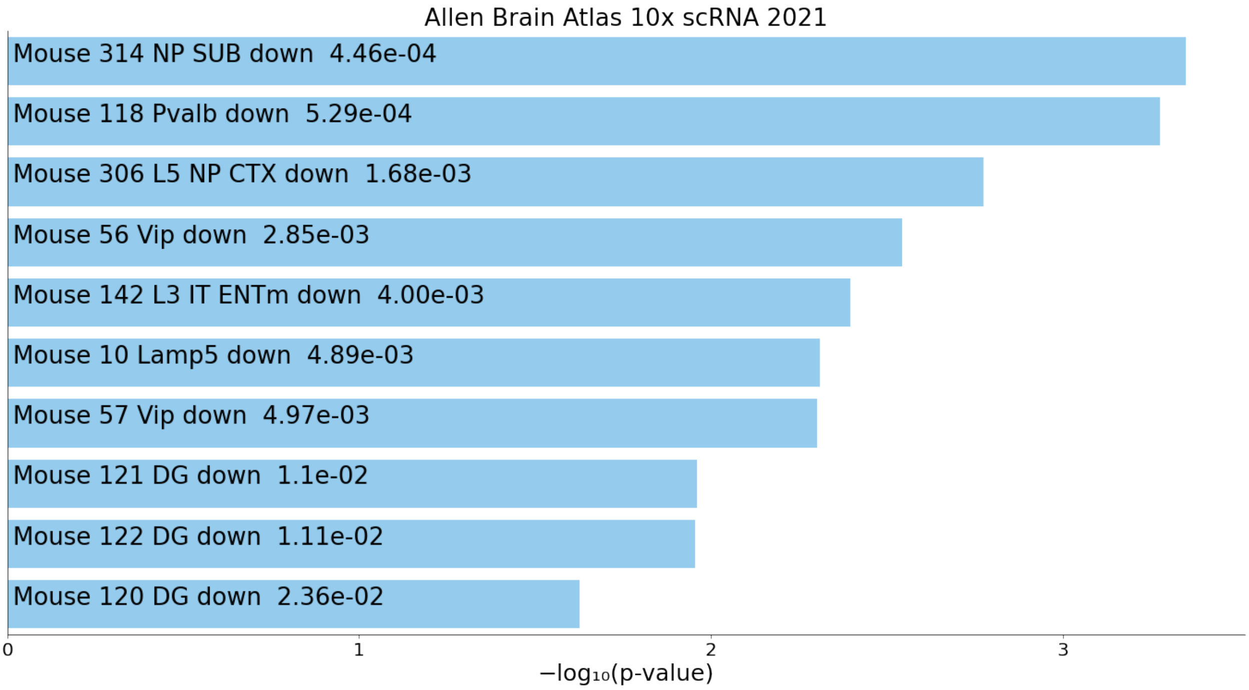


Figure S39 hippocampal_layer_down2(brain up)

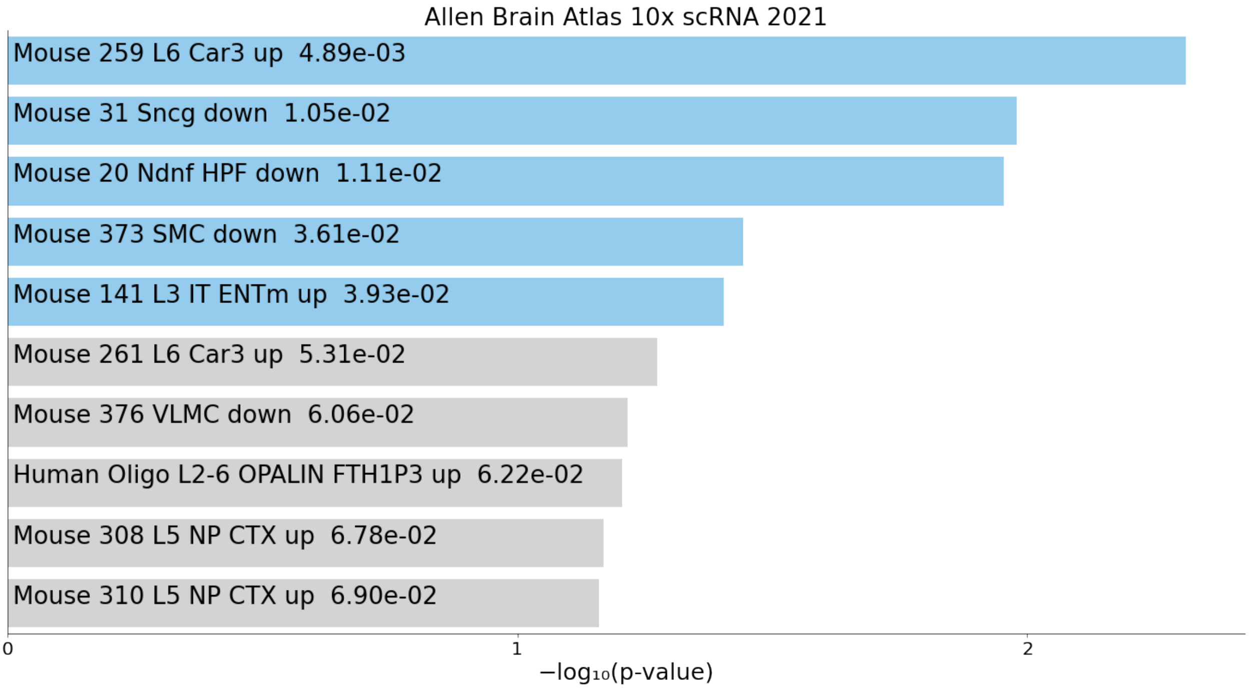


Figure S40 frontal_cortex_up2

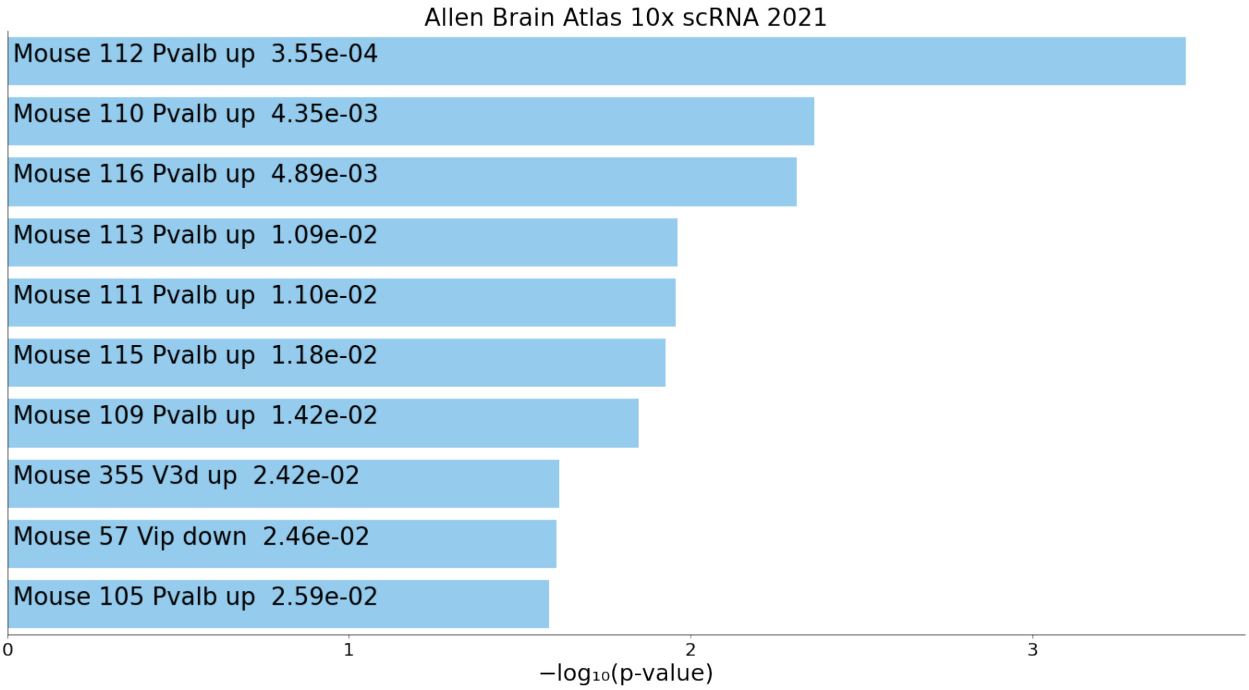


Figure S41 frontal_cortex_down2(brain up)
