## Supplementary material for "Somatostatin-expressing neurons regulate sleep deprivation and recovery": Suuplementary_Tables.pdf

### Table alpha genome enrichr

Kobaysahi Kenta

September 2025

#### $\ell_1 = 2$

Table S1: left cerebral cortex up1

| Index | Name | P-value | Adjusted p-value | Odds Ratio | Combined score |
|---|---|---|---|---|---|
| 1 | Mouse 96 Sst up | 0.0004666 | 0.08772 | 2.38 | 18.26 |
| 2 | Mouse 312 NP SUB up | 0.001048 | 0.09848 | 16.60 | 113.92 |
| 3 | Mouse 309 L5 NP CTX up | 0.02963 | 0.55990 | 40.19 | 141.43 |
| 4 | Mouse 318 CT SUB up | 0.02963 | 0.55990 | 40.19 | 141.43 |
| 5 | Mouse 314 NP SUB up | 0.03931 | 0.55990 | 28.71 | 92.90 |
| 6 | Mouse 152 L2 IT ENTm down | 0.03931 | 0.55990 | 28.71 | 92.90 |
| 7 | Mouse 165 L2/3 IT ENTl down | 0.03931 | 0.55990 | 28.71 | 92.90 |
| 8 | Human Inh L2-5 VIP BSPRY down | 0.04317 | 0.55990 | 6.33 | 19.88 |
| 9 | Mouse 148 L2 IT PAR down | 0.04412 | 0.55990 | 25.12 | 78.38 |
| 10 | Mouse 293 CA3 down | 0.05366 | 0.55990 | 20.09 | 58.77 |

Table S2: left cerebral cortex down1(brain up)

| Index | Name | P-value | Adjusted p-value | Odds Ratio | Combined score |
|---|---|---|---|---|---|
| 1 | Mouse 33 Sncg down | 0.00004230 | 0.01320 | 23.65 | 238.16 |
| 2 | Human Astro L1-6 FGFR3 AQP1 down | 0.0002117 | 0.02536 | 3.77 | 31.88 |
| 3 | Mouse 359 OPC down | 0.0002833 | 0.02536 | 3.44 | 28.12 |
| 4 | Mouse 19 Pax6 down | 0.0003251 | 0.02536 | 25.61 | 205.71 |
| 5 | Human OPC L1-6 PDGFRA COL20A1 down | 0.0004251 | 0.02652 | 3.01 | 23.37 |
| 6 | Mouse 100 Sst down | 0.0005934 | 0.03086 | 20.48 | 152.19 |
| 7 | Human Inh L1-2 VIP SCML4 down | 0.0008709 | 0.03882 | 58.00 | 408.65 |
| 8 | Human Astro L1-6 FGFR3 PLCG1 down | 0.001462 | 0.05094 | 2.99 | 19.52 |
| 9 | Mouse 318 CT SUB down | 0.001598 | 0.05094 | 8.60 | 55.35 |
| 10 | Mouse 22 Ndnf HPF down | 0.001862 | 0.05094 | 36.90 | 231.95 |

Table S3: right cerebral cortex up1

| Index | Name | P-value | Adjusted p-value | Odds Ratio | Combined score |
|---|---|---|---|---|---|
| 1 | Mouse 96 Sst up | 0.0001824 | 0.03374 | 2.51 | 21.65 |
| 2 | Mouse 312 NP SUB up | 0.01706 | 0.64240 | 10.67 | 43.42 |
| 3 | Human Inh L5-6 PVALB ZFPM2-AS1 up | 0.02236 | 0.64240 | 5.23 | 19.88 |
| 4 | Mouse 309 L5 NP CTX up | 0.02963 | 0.64240 | 40.19 | 141.43 |
| 5 | Mouse 318 CT SUB up | 0.02963 | 0.64240 | 40.19 | 141.43 |
| 6 | Mouse 97 Sst up | 0.03426 | 0.64240 | 1.97 | 6.66 |
| 7 | Mouse 314 NP SUB up | 0.03931 | 0.64240 | 28.71 | 92.90 |
| 8 | Human Inh L3-5 SST CDH3 down | 0.05686 | 0.64240 | 5.39 | 15.47 |
| 9 | Mouse 208 L5 IT CTX up | 0.05839 | 0.64240 | 18.26 | 51.88 |
| 10 | Mouse 132 L2/3 IT PPP down | 0.05915 | 0.64240 | 3.51 | 9.91 |

Table S4: right cerebral cortex down1(brain up)

| Index | Name | P-value | Adjusted p-value | Odds Ratio | Combined score |
|---|---|---|---|---|---|
| 1 | Mouse 123 DG down | 0.000005007 | 0.001863 | 23.22 | 283.42 |
| 2 | Mouse 293 CA3 down | 0.000019440 | 0.003615 | 76.90 | 834.26 |
| 3 | Mouse 362 Oligo down | 0.000038480 | 0.004772 | 4.01 | 40.73 |
| 4 | Mouse 57 Vip down | 0.000119200 | 0.011090 | 6.96 | 62.85 |
| 5 | Mouse 40 Vip down | 0.000231400 | 0.014930 | 14.76 | 123.60 |
| 6 | Mouse 119 Pvalb Vipr2 down | 0.000279500 | 0.014930 | 14.01 | 114.65 |
| 7 | Mouse 54 Vip down | 0.000280900 | 0.014930 | 7.28 | 59.52 |
| 8 | Mouse 56 Vip down | 0.000419900 | 0.019530 | 6.73 | 52.32 |
| 9 | Human Inh L5-6 PVALB FAM150B down | 0.000492400 | 0.020010 | 21.95 | 167.17 |
| 10 | Mouse 363 Oligo down | 0.000538000 | 0.020010 | 3.38 | 25.43 |
| 11 | Human Inh L6 SST TH down | 0.000768200 | 0.021600 | 18.62 | 133.53 |
| 12 | Mouse 103 Sst up | 0.000768200 | 0.021600 | 18.62 | 133.53 |
| 13 | Mouse 373 SMC down | 0.000770800 | 0.021600 | 4.00 | 28.68 |
| 14 | Mouse 112 Pvalb down | 0.000832900 | 0.021600 | 18.07 | 128.13 |
| 15 | Mouse 71 Sst down | 0.000870900 | 0.021600 | 58.00 | 408.65 |
| 16 | Mouse 33 Sncg down | 0.000972600 | 0.021650 | 17.07 | 118.36 |
| 17 | Mouse 356 Astro down | 0.001040000 | 0.021650 | 3.53 | 24.26 |
| 18 | Mouse 104 Sst down | 0.001048000 | 0.021650 | 16.60 | 113.92 |
| 19 | Human VLMC L1-5 PDGFRA COLEC12 down | 0.001199000 | 0.023480 | 3.24 | 21.80 |
| 20 | Mouse 103 Sst down | 0.001478000 | 0.027490 | 14.62 | 95.30 |
| 21 | Mouse 1 CR down | 0.001594000 | 0.028240 | 3.12 | 20.10 |
| 22 | Mouse 55 Vip down | 0.001846000 | 0.028770 | 8.25 | 51.93 |
| 23 | Mouse 22 Ndnf HPF down | 0.001862000 | 0.028770 | 36.90 | 231.95 |
| 24 | Mouse 300 CA3 down | 0.001862000 | 0.028770 | 36.90 | 231.95 |
| 25 | Human Inh L3-5 VIP IGDCC3 down | 0.002122000 | 0.028770 | 12.79 | 78.73 |

Table S5: hippocampal layer up1

| Index | Name | P-value | Adjusted p-value | Odds Ratio | Combined score |
|---|---|---|---|---|---|
| 1 | Mouse 96 Sst up | 7.990e-7 | 0.000142 | 3.24 | 45.51 |
| 2 | Mouse 97 Sst up | 0.006042 | 0.537700 | 2.39 | 12.21 |
| 3 | Mouse 310 L5 NP CTX up | 0.009197 | 0.545700 | 7.38 | 34.62 |
| 4 | Mouse 371 Peri up | 0.020430 | 0.663900 | 9.65 | 37.54 |
| 5 | Mouse 19 Pax6 up | 0.021310 | 0.663900 | 9.42 | 36.27 |
| 6 | Mouse 309 L5 NP CTX up | 0.029630 | 0.663900 | 40.19 | 141.43 |
| 7 | Mouse 318 CT SUB up | 0.029630 | 0.663900 | 40.19 | 141.43 |
| 8 | Mouse 70 Sst up | 0.033060 | 0.663900 | 7.36 | 25.11 |
| 9 | Mouse 314 NP SUB up | 0.039310 | 0.663900 | 28.71 | 92.90 |
| 10 | Human Inh L5-6 SST FBN2 down | 0.055560 | 0.663900 | 5.47 | 15.80 |

Table S6: hippocampal layer down1(brain up)

| Index | Name | P-value | Adjusted p-value | Odds Ratio | Combined score |
|---|---|---|---|---|---|
| 1 | Mouse 123 DG down | 0.000005007 | 0.001778 | 23.22 | 283.42 |
| 2 | Mouse 56 Vip down | 0.000052470 | 0.009314 | 7.98 | 78.62 |
| 3 | Mouse 57 Vip down | 0.000119200 | 0.014110 | 6.96 | 62.85 |
| 4 | Mouse 119 Pvalb Vipr2 down | 0.000279500 | 0.023340 | 14.01 | 114.65 |
| 5 | Human VLMC L1-5 PDGFRA COLEC12 down | 0.000328800 | 0.023340 | 3.58 | 28.72 |
| 6 | Human Inh L5-6 PVALB FAM150B down | 0.000492400 | 0.026330 | 21.95 | 167.17 |
| 7 | Mouse 362 Oligo down | 0.000582000 | 0.026330 | 3.35 | 24.93 |
| 8 | Mouse 100 Sst down | 0.000593400 | 0.026330 | 20.48 | 152.19 |
| 9 | Human Inh L6 SST TH down | 0.000768200 | 0.026880 | 18.62 | 133.53 |
| 10 | Mouse 373 SMC down | 0.000770800 | 0.026880 | 4.00 | 28.68 |

Table S7: frontal cortex up1

| Index | Name | P-value | Adjusted p-value | Odds Ratio | Combined score |
|---|---|---|---|---|---|
| 1 | Mouse 188 L4/5 IT CTX down | 0.001295 | 0.2450 | 15.36 | 102.11 |
| 2 | Mouse 97 Sst up | 0.002250 | 0.2450 | 2.61 | 15.89 |
| 3 | Mouse 118 Pvalb down | 0.004361 | 0.2450 | 6.44 | 34.98 |
| 4 | Mouse 43 Vip down | 0.004974 | 0.2450 | 6.19 | 32.84 |
| 5 | Mouse 356 Astro up | 0.005635 | 0.2450 | 8.89 | 46.03 |
| 6 | Mouse 44 Vip down | 0.006520 | 0.2450 | 5.72 | 28.77 |
| 7 | Mouse 127 L2/3 IT APr down | 0.008008 | 0.2450 | 16.22 | 78.32 |
| 8 | Mouse 347 L6b CTX up | 0.010100 | 0.2450 | 7.13 | 32.75 |
| 9 | Mouse 102 Sst down | 0.010470 | 0.2450 | 13.98 | 63.75 |
| 10 | Mouse 42 Vip down | 0.011050 | 0.2450 | 6.88 | 31.02 |

Table S8: frontal cortex down1(brain up)

| Index | Name | P-value | Adjusted p-value | Odds Ratio | Combined score |
|---|---|---|---|---|---|
| 1 | Mouse 373 SMC down | 0.00003329 | 0.007464 | 5.02 | 51.78 |
| 2 | Mouse 370 Endo down | 0.00005046 | 0.007464 | 4.42 | 43.78 |
| 3 | Mouse 123 DG down | 0.00011360 | 0.007464 | 17.98 | 163.35 |
| 4 | Mouse 57 Vip down | 0.00011920 | 0.007464 | 6.96 | 62.85 |
| 5 | Mouse 363 Oligo down | 0.00014290 | 0.007464 | 3.71 | 32.82 |
| 6 | Mouse 355 V3d down | 0.00014340 | 0.007464 | 4.23 | 37.43 |
| 7 | Mouse 362 Oligo down | 0.00015600 | 0.007464 | 3.67 | 32.19 |
| 8 | Mouse 360 OPC down | 0.00023050 | 0.008104 | 3.35 | 28.03 |
| 9 | Mouse 374 VLMC down | 0.00024790 | 0.008104 | 4.28 | 35.57 |
| 10 | Mouse 356 Astro down | 0.00025970 | 0.008104 | 3.94 | 32.49 |
| 11 | Mouse 376 VLMC down | 0.00026610 | 0.008104 | 4.67 | 38.41 |
| 12 | Human Inh L5-6 PVALB FAM150B down | 0.00049240 | 0.012700 | 21.95 | 167.17 |
| 13 | Mouse 378 Micro down | 0.00049280 | 0.012700 | 4.76 | 36.25 |
| 14 | Mouse 361 Oligo down | 0.00061730 | 0.014770 | 3.15 | 23.31 |
| 15 | Human Inh L6 SST TH down | 0.00076820 | 0.017160 | 18.62 | 133.53 |
| 16 | Mouse 6 Lamp5 Lhx6 down | 0.00099360 | 0.020800 | 4.82 | 33.32 |
| 17 | Human VLMC L1-5 PDGFRA COLEC12 down | 0.00119900 | 0.022420 | 3.24 | 21.80 |
| 18 | Mouse 371 Peri down | 0.00120500 | 0.022420 | 4.12 | 27.72 |
| 19 | Mouse 293 CA3 down | 0.00132200 | 0.023310 | 45.10 | 298.98 |
| 20 | Mouse 358 Astro down | 0.00164200 | 0.027500 | 3.31 | 21.23 |
| 21 | Mouse 366 Oligo down | 0.00207000 | 0.032980 | 3.01 | 18.62 |
| 22 | Mouse 76 Sst down | 0.00216600 | 0.032980 | 33.82 | 207.50 |
| 23 | Mouse 377 Micro down | 0.00238500 | 0.034740 | 4.11 | 24.81 |
| 24 | Mouse 121 DG down | 0.00265100 | 0.037010 | 4.03 | 23.90 |
| 25 | Mouse 56 Vip down | 0.00285300 | 0.038230 | 5.52 | 32.33 |

$\ell_1 = 3$

Table S9: left cerebral cortex up2

| Index | Name | P-value | Adjusted p-value | Odds Ratio | Combined score |
|---|---|---|---|---|---|
| 1 | Mouse 374 VLMC up | 0.02405 | 0.8553 | 8.81 | 32.83 |
| 2 | Mouse 70 Sst up | 0.03306 | 0.8553 | 7.36 | 25.11 |
| 3 | Human Inh L5-6 PVALB SST CRHR2 down | 0.04412 | 0.8553 | 25.12 | 78.38 |
| 4 | Human Inh L3-5 SST CDH3 down | 0.05686 | 0.8553 | 5.39 | 15.47 |
| 5 | Mouse 293 CA3 up | 0.05820 | 0.8553 | 2.49 | 7.08 |
| 6 | Mouse 15 Lamp5 up | 0.05839 | 0.8553 | 18.26 | 51.88 |
| 7 | Mouse 28 Sncg up | 0.06625 | 0.8553 | 2.72 | 7.39 |
| 8 | Mouse 55 Vip up | 0.06779 | 0.8553 | 15.45 | 41.59 |
| 9 | Mouse 308 L5 NP CTX up | 0.06779 | 0.8553 | 15.45 | 41.59 |
| 10 | Mouse 30 Sncg up | 0.07366 | 0.8553 | 1.99 | 5.20 |

Table S10: left cerebral cortex down2 (brain up)

| Index | Name | P-value | Adjusted p-value | Odds Ratio | Combined score |
|---|---|---|---|---|---|
| 1 | Mouse 236 L3 RSP-ACA down | 0.0006796 | 0.14026 | 67.67 | 493.57 |
| 2 | Human Astro L1 FGFR3 SERPINI2 down | 0.0011400 | 0.14022 | 2.94 | 19.91 |
| 3 | Mouse 90 Sst down | 0.0032050 | 0.19350 | 27.05 | 155.37 |
| 4 | Mouse 267 SUB-ProS down | 0.0035940 | 0.19350 | 25.36 | 142.75 |
| 5 | Mouse 91 Sst down | 0.0044340 | 0.19350 | 22.54 | 122.14 |
| 6 | Mouse 379 PVM down | 0.0051440 | 0.19350 | 3.55 | 18.72 |
| 7 | Human Inh L3-6 PAX6 LINC01497 down | 0.0068890 | 0.19350 | 17.64 | 87.79 |
| 8 | Mouse 66 Sst down | 0.0085960 | 0.19350 | 15.60 | 74.20 |
| 9 | Mouse 46 Vip down | 0.0089080 | 0.19350 | 7.47 | 35.29 |
| 10 | Mouse 359 OPC down | 0.0093620 | 0.19350 | 2.55 | 11.91 |

Table S11: right cerebral cortex up2

| Index | Name | P-value | Adjusted p-value | Odds Ratio | Combined score |
|---|---|---|---|---|---|
| 1 | Mouse 96 Sst up | 0.005729 | 0.7639 | 2.00 | 10.33 |
| 2 | Mouse 309 L5 NP CTX up | 0.029630 | 0.7639 | 40.19 | 141.43 |
| 3 | Mouse 318 CT SUB up | 0.029630 | 0.7639 | 40.19 | 141.43 |
| 4 | Mouse 314 NP SUB up | 0.039310 | 0.7639 | 28.71 | 92.90 |
| 5 | Mouse 378 Micro up | 0.050460 | 0.7639 | 5.78 | 17.27 |
| 6 | Mouse 208 L5 IT CTX up | 0.058390 | 0.7639 | 18.26 | 51.88 |
| 7 | Mouse 89 Sst up | 0.060820 | 0.7639 | 5.19 | 14.52 |
| 8 | Mouse 313 NP SUB up | 0.063100 | 0.7639 | 16.74 | 46.25 |
| 9 | Mouse 306 L5 NP CTX up | 0.063100 | 0.7639 | 16.74 | 46.25 |
| 10 | Mouse 28 Sncg up | 0.066250 | 0.7639 | 2.72 | 7.39 |

Table S12: right cerebral cortex down2 (brain up)

| Index | Name | P-value | Adjusted p-value | Odds Ratio | Combined score |
|---|---|---|---|---|---|
| 1 | Mouse 46 Vip down | 0.00006752 | 0.02093 | 13.04 | 125.22 |
| 2 | Human Inh L5-6 PVALB FAM150B down | 0.00049240 | 0.05025 | 21.95 | 167.17 |
| 3 | Mouse 118 Pvalb down | 0.00052920 | 0.05025 | 8.19 | 61.82 |
| 4 | Mouse 103 Sst up | 0.00076820 | 0.05025 | 18.62 | 133.53 |
| 5 | Mouse 57 Vip down | 0.00083500 | 0.05025 | 5.87 | 41.62 |
| 6 | Mouse 33 Sncg down | 0.00097260 | 0.05025 | 17.07 | 118.36 |
| 7 | Mouse 231 L6 IT CTX down | 0.00120900 | 0.05352 | 15.75 | 105.81 |
| 8 | Mouse 114 Pvalb down | 0.00147800 | 0.05595 | 14.62 | 95.30 |
| 9 | Mouse 306 L5 NP CTX down | 0.00167700 | 0.05595 | 13.96 | 89.20 |
| 10 | Mouse 54 Vip down | 0.00206000 | 0.05595 | 5.97 | 36.91 |

Table S13: hippocampal layer up2

| Index | Name | P-value | Adjusted p-value | Odds Ratio | Combined score |
|---|---|---|---|---|---|
| 1 | Mouse 96 Sst up | 0.001134 | 0.1679 | 2.25 | 15.27 |
| 2 | Mouse 30 Sncg up | 0.010960 | 0.7873 | 2.63 | 11.86 |
| 3 | Mouse 371 Peri up | 0.020430 | 0.7873 | 9.65 | 37.54 |
| 4 | Human Exc L6 FEZF2 PDYN up | 0.021830 | 0.7873 | 2.13 | 8.14 |
| 5 | Mouse 97 Sst up | 0.034260 | 0.7873 | 1.97 | 6.66 |
| 6 | Mouse 374 VLMC down | 0.043830 | 0.7873 | 2.44 | 7.64 |
| 7 | Mouse 229 L6 IT CTX up | 0.052560 | 0.7873 | 2.95 | 8.70 |
| 8 | Mouse 231 L6 IT CTX up | 0.052990 | 0.7873 | 5.62 | 16.51 |
| 9 | Human Exc L3 RORB OTOGL up | 0.077100 | 0.7873 | 13.39 | 34.32 |
| 10 | Mouse 31 Sncg up | 0.083340 | 0.7873 | 2.05 | 5.10 |

Table S14: hippocampal layer down2(brain up)

| Index | Name | P-value | Adjusted p-value | Odds Ratio | Combined score |
|---|---|---|---|---|---|
| 1 | Mouse 306 L5 NP CTX down | 0.00008897 | 0.01657 | 19.24 | 179.47 |
| 2 | Mouse 57 Vip down | 0.00011920 | 0.01657 | 6.96 | 62.85 |
| 3 | Human Inh L5-6 PVALB FAM150B down | 0.00049240 | 0.03160 | 21.95 | 167.17 |
| 4 | Mouse 118 Pvalb down | 0.00052920 | 0.03160 | 8.19 | 61.82 |
| 5 | Mouse 122 DG down | 0.00056830 | 0.03160 | 4.65 | 34.78 |
| 6 | Mouse 309 L5 NP CTX down | 0.00212200 | 0.08601 | 12.79 | 78.73 |
| 7 | Mouse 76 Sst down | 0.00216600 | 0.08601 | 33.82 | 207.50 |
| 8 | Mouse 121 DG down | 0.00265100 | 0.08813 | 4.03 | 23.90 |
| 9 | Mouse 56 Vip down | 0.00285300 | 0.08813 | 5.52 | 32.33 |
| 10 | Human Inh L1-2 SST CLIC6 down | 0.00584800 | 0.14800 | 19.32 | 99.33 |

Table S15: frontal cortex up2

| Index | Name | P-value | Adjusted p-value | Odds Ratio | Combined score |
|---|---|---|---|---|---|
| 1 | Mouse 259 L6 Car3 up | 0.0001511 | 0.02659 | 34.16 | 300.54 |
| 2 | Mouse 158 L2/3 IT ENTl up | 0.0040040 | 0.24800 | 23.87 | 131.77 |
| 3 | Mouse 356 Astro up | 0.0056350 | 0.24800 | 8.89 | 46.03 |
| 4 | Mouse 378 Micro up | 0.0056350 | 0.24800 | 8.89 | 46.03 |
| 5 | Mouse 310 L5 NP CTX up | 0.0091970 | 0.32370 | 7.38 | 34.62 |
| 6 | Mouse 377 Micro up | 0.0130900 | 0.38410 | 6.45 | 27.95 |
| 7 | Mouse 261 L6 Car3 up | 0.0162700 | 0.40900 | 3.11 | 12.82 |
| 8 | Mouse 258 L6 Car3 up | 0.0299300 | 0.62320 | 7.79 | 27.33 |
| 9 | Mouse 373 SMC down | 0.0360700 | 0.62320 | 2.57 | 8.53 |
| 10 | Mouse 141 L3 IT ENTm up | 0.0393100 | 0.62320 | 28.71 | 92.90 |
| 11 | Human Exc L3-5 RORB TNNT2 up | 0.0467600 | 0.62320 | 6.04 | 18.50 |
| 12 | Mouse 260 L6 Car3 up | 0.0591500 | 0.62320 | 3.51 | 9.91 |
| 13 | Mouse 376 VLMC down | 0.0606300 | 0.62320 | 2.46 | 6.89 |
| 14 | Mouse 379 PVM up | 0.0648800 | 0.62320 | 4.99 | 13.66 |
| 15 | Mouse 308 L5 NP CTX up | 0.0677900 | 0.62320 | 15.45 | 41.59 |
| 16 | Mouse 97 Sst up | 0.0718300 | 0.62320 | 1.77 | 4.67 |
| 17 | Mouse 142 L3 IT ENTm up | 0.0724600 | 0.62320 | 14.35 | 37.66 |
| 18 | Mouse 347 L6b CTX up | 0.0732500 | 0.62320 | 4.65 | 12.15 |
| 19 | Human Exc L3 RORB OTOGL up | 0.0771000 | 0.62320 | 13.39 | 34.32 |
| 20 | Mouse 262 L6 Car3 up | 0.0798300 | 0.62320 | 3.08 | 7.78 |
| 21 | Mouse 15 Lamp5 down | 0.0817100 | 0.62320 | 12.55 | 31.44 |
| 22 | Mouse 37 Sncg down | 0.0863100 | 0.62320 | 11.81 | 28.94 |
| 23 | Mouse 358 Astro up | 0.0894400 | 0.62320 | 4.12 | 9.96 |
| 24 | Mouse 185 L2 IT RSP-ACA down | 0.0971100 | 0.62320 | 2.82 | 6.57 |
| 25 | Mouse 329 L6 CT CTX up | 0.1045000 | 0.62320 | 9.56 | 21.60 |

Table S16: frontal cortex down2(brain up)

| Index | Name | P-value | Adjusted p-value | Odds Ratio | Combined score |
|---|---|---|---|---|---|
| 1 | Human Inh L5-6 PVALB FAM150B down | 0.0004924 | 0.08099 | 21.95 | 167.17 |
| 2 | Mouse 103 Sst up | 0.0007682 | 0.08099 | 18.62 | 133.53 |
| 3 | Mouse 57 Vip down | 0.0008350 | 0.08099 | 5.87 | 41.62 |
| 4 | Mouse 376 VLMC down | 0.0012610 | 0.08933 | 4.09 | 27.32 |
| 5 | Mouse 113 Pvalb up | 0.0017300 | 0.08933 | 6.22 | 39.55 |
| 6 | Mouse 76 Sst down | 0.0021660 | 0.08933 | 33.82 | 207.50 |
| 7 | Mouse 378 Micro down | 0.0023850 | 0.08933 | 4.11 | 24.81 |
| 8 | Mouse 73 Sst up | 0.0028370 | 0.08933 | 28.99 | 170.01 |
| 9 | Mouse 70 Sst up | 0.0029180 | 0.08933 | 11.37 | 66.34 |
| 10 | Mouse 112 Pvalb up | 0.0031970 | 0.08933 | 7.05 | 40.48 |

#### $\ell_1 = 2$ and $\ell_1 = 3$

Table S17: left cerebral cortex up

| Index | Name | P-value | Adjusted p-value | Odds Ratio | Combined score |
|---|---|---|---|---|---|
| 1 | Mouse 96 Sst up | 7.990e-7 | 0.000172 | 3.24 | 45.51 |
| 2 | Mouse 129 L2/3 IT APr down | 0.002728 | 0.293300 | 5.58 | 32.94 |
| 3 | Human Inh L5-6 PVALB FAM150B down | 0.010470 | 0.469600 | 13.98 | 63.75 |
| 4 | Mouse 132 L2/3 IT PPP down | 0.012040 | 0.469600 | 4.75 | 21.00 |
| 5 | Mouse 125 DG down | 0.019890 | 0.469600 | 2.97 | 11.63 |
| 6 | Human Exc L5 THEMIS LINC01116 up | 0.019900 | 0.469600 | 1.80 | 7.04 |
| 7 | Mouse 120 DG down | 0.023570 | 0.469600 | 2.85 | 10.68 |
| 8 | Mouse 57 Vip down | 0.024620 | 0.469600 | 3.80 | 14.06 |
| 9 | Mouse 53 Vip down | 0.026410 | 0.469600 | 4.89 | 17.78 |
| 10 | Mouse 51 Vip down | 0.028580 | 0.469600 | 4.74 | 16.85 |

Table S18: left cerebral cortex down(brain up)

| Index | Name | P-value | Adjusted p-value | Odds Ratio | Combined score |
|---|---|---|---|---|---|
| 1 | Mouse 379 PVM down | 0.0002570 | 0.03358 | 4.69 | 38.77 |
| 2 | Mouse 359 OPC down | 0.0002833 | 0.03358 | 3.44 | 28.12 |
| 3 | Mouse 361 Oligo down | 0.0006173 | 0.03734 | 3.15 | 23.31 |
| 4 | Mouse 360 OPC down | 0.0007853 | 0.03734 | 3.07 | 21.94 |
| 5 | Mouse 356 Astro down | 0.0010400 | 0.03734 | 3.53 | 24.26 |
| 6 | Mouse 374 VLMC down | 0.0010790 | 0.03734 | 3.81 | 26.00 |
| 7 | Mouse 96 Sst up | 0.0011340 | 0.03734 | 2.25 | 15.27 |
| 8 | Mouse 376 VLMC down | 0.0012610 | 0.03734 | 4.09 | 27.32 |
| 9 | Mouse 363 Oligo down | 0.0018560 | 0.04675 | 3.06 | 19.23 |
| 10 | Human Oligo L2-6 OPALIN FTH1P3 down | 0.0019730 | 0.04675 | 2.88 | 17.93 |

Table S19: right cerebral cortex up

| Index | Name | P-value | Adjusted p-value | Odds Ratio | Combined score |
|---|---|---|---|---|---|
| 1 | Mouse 96 Sst up | 7.990e-7 | 0.000169 | 3.24 | 45.51 |
| 2 | Human Exc L5 THEMIS LINC01116 up | 0.009972 | 0.539400 | 1.92 | 8.84 |
| 3 | Mouse 6 Lamp5 Lhx6 down | 0.020500 | 0.539400 | 3.35 | 13.02 |
| 4 | Human Exc L5 THEMIS RGPD6 up | 0.021510 | 0.539400 | 3.30 | 12.69 |
| 5 | Human Exc L6 FEZF2 PDYN up | 0.021830 | 0.539400 | 2.13 | 8.14 |
| 6 | Mouse 112 Pvalb up | 0.022840 | 0.539400 | 5.18 | 19.59 |
| 7 | Mouse 309 L5 NP CTX up | 0.029630 | 0.539400 | 40.19 | 141.43 |
| 8 | Mouse 318 CT SUB up | 0.029630 | 0.539400 | 40.19 | 141.43 |
| 9 | Mouse 124 DG down | 0.030360 | 0.539400 | 3.55 | 12.40 |
| 10 | Mouse 97 Sst up | 0.034260 | 0.539400 | 1.97 | 6.66 |

Table S20: right cerebral cortex down(brain up)

| **Index** | **Name** | **P-value** | **Adjusted p-value** | **Odds Ratio** | **Combined score** |
|---|---|---|---|---|---|
| 1 | Mouse 267 SUB-ProS down | 0.00009371 | 0.01885 | 41.00 | 380.29 |
| 2 | Mouse 123 DG down | 0.00011360 | 0.01885 | 17.98 | 163.35 |
| 3 | Human Inh L5-6 PVALB FAM150B down | 0.00049240 | 0.05450 | 21.95 | 167.17 |
| 4 | Human Inh L3-6 VIP ZIM2-AS1 down | 0.00083290 | 0.06537 | 18.07 | 128.13 |
| 5 | Mouse 376 VLMC down | 0.00126100 | 0.06537 | 4.09 | 27.32 |
| 6 | Mouse 364 Oligo down | 0.00153400 | 0.06537 | 3.34 | 21.67 |
| 7 | Mouse 22 Ndnf HPF down | 0.00186200 | 0.06537 | 36.90 | 231.95 |
| 8 | Mouse 292 CA3 down | 0.00186200 | 0.06537 | 36.90 | 231.95 |
| 9 | Mouse 294 CA3 down | 0.00186200 | 0.06537 | 36.90 | 231.95 |
| 10 | Mouse 76 Sst down | 0.00216600 | 0.06537 | 33.82 | 207.50 |

Table S21: hippocampal layer up

| **Index** | **Name** | **P-value** | **Adjusted p-value** | **Odds Ratio** | **Combined score** |
|---|---|---|---|---|---|
| 1 | Mouse 96 Sst up | 0.000002603 | 0.0005441 | 3.09 | 39.70 |
| 2 | Mouse 97 Sst up | 0.000074880 | 0.0078250 | 3.29 | 31.23 |
| 3 | Human Exc L5 THEMIS LINC01116 up | 0.004726000 | 0.3293000 | 2.04 | 10.92 |
| 4 | Human Exc L5-6 FEZF2 LPO up | 0.006643000 | 0.3471000 | 2.69 | 13.49 |
| 5 | Mouse 358 Astro up | 0.013820000 | 0.4508000 | 6.31 | 27.03 |
| 6 | Mouse 371 Peri up | 0.020430000 | 0.4508000 | 9.65 | 37.54 |
| 7 | Human Exc L5 THEMIS RGPD6 up | 0.021510000 | 0.4508000 | 3.30 | 12.69 |
| 8 | Mouse 309 L5 NP CTX up | 0.029630000 | 0.4508000 | 40.19 | 141.43 |
| 9 | Mouse 318 CT SUB up | 0.029630000 | 0.4508000 | 40.19 | 141.43 |
| 10 | Human Exc L5-6 FEZF2 OR1L8 up | 0.034480000 | 0.4508000 | 2.13 | 7.17 |

Table S22: hippocampal layer down(brain up)

| Index | Name | P-value | Adjusted p-value | Odds Ratio | Combined score |
|---|---|---|---|---|---|
| 1 | Mouse 376 VLMC down | 0.00005016 | 0.008295 | 5.26 | 52.03 |
| 2 | Mouse 374 VLMC down | 0.00005136 | 0.008295 | 4.78 | 47.17 |
| 3 | Mouse 378 Micro down | 0.00008985 | 0.009171 | 5.43 | 50.59 |
| 4 | Mouse 123 DG down | 0.00011360 | 0.009171 | 17.98 | 163.35 |
| 5 | Human VLMC L1-5 PDGFRA COLEC12 down | 0.00032880 | 0.017690 | 3.58 | 28.72 |
| 6 | Human Inh L2-5 VIP BSPRY down | 0.00033430 | 0.017690 | 13.33 | 106.70 |
| 7 | Human Inh L3-6 VIP UG0898H09 down | 0.00044640 | 0.017690 | 22.76 | 175.61 |
| 8 | Human Inh L5-6 PVALB FAM150B down | 0.00049240 | 0.017690 | 21.95 | 167.17 |
| 9 | Mouse 377 Micro down | 0.00049280 | 0.017690 | 4.76 | 36.25 |
| 10 | Human Oligo L2-6 OPALIN FTH1P3 down | 0.00061100 | 0.019740 | 3.16 | 23.37 |
| 11 | Mouse 373 SMC down | 0.00077080 | 0.022420 | 4.00 | 28.68 |
| 12 | Human Inh L3-6 VIP ZIM2-AS1 down | 0.00083290 | 0.022420 | 18.07 | 128.13 |
| 13 | Mouse 370 Endo down | 0.00091880 | 0.022830 | 3.59 | 25.12 |
| 14 | Mouse 371 Peri down | 0.00120500 | 0.026340 | 4.12 | 27.72 |
| 15 | Mouse 379 PVM down | 0.00122300 | 0.026340 | 4.11 | 27.58 |
| 16 | Human Astro L1-6 FGFR3 PLCG1 down | 0.00146200 | 0.026910 | 2.99 | 19.52 |
| 17 | Mouse 364 Oligo down | 0.00153400 | 0.026910 | 3.34 | 21.67 |
| 18 | Mouse 363 Oligo down | 0.00185600 | 0.026910 | 3.06 | 19.23 |
| 19 | Mouse 22 Ndnf HPF down | 0.00186200 | 0.026910 | 36.90 | 231.95 |
| 20 | Mouse 292 CA3 down | 0.00186200 | 0.026910 | 36.90 | 231.95 |
| 21 | Mouse 294 CA3 down | 0.00186200 | 0.026910 | 36.90 | 231.95 |
| 22 | Mouse 361 Oligo down | 0.00199100 | 0.026910 | 2.88 | 17.88 |
| 23 | Mouse 366 Oligo down | 0.00207000 | 0.026910 | 3.01 | 18.62 |
| 24 | Human Inh L3-5 VIP IGDCC3 down | 0.00212200 | 0.026910 | 12.79 | 78.73 |
| 25 | Mouse 76 Sst down | 0.00216600 | 0.026910 | 33.82 | 207.50 |

Table S23: frontal cortex up

| Index | Name | P-value | Adjusted p-value | Odds Ratio | Combined score |
|---|---|---|---|---|---|
| 1 | Mouse 96 Sst up | 2.343e-7 | 0.000057 | 3.40 | 51.90 |
| 2 | Human Inh L1 PAX6 MIR101-1 up | 0.001384 | 0.167500 | 14.98 | 98.61 |
| 3 | Mouse 301 CA2-IG-FC up | 0.003704 | 0.242200 | 10.40 | 58.23 |
| 4 | Mouse 158 L2/3 IT ENTl up | 0.004004 | 0.242200 | 23.87 | 131.77 |
| 5 | Mouse 378 Micro up | 0.005635 | 0.246200 | 8.89 | 46.03 |
| 6 | Human Inh L2 PAX6 FREM2 up | 0.006103 | 0.246200 | 3.88 | 19.79 |
| 7 | Mouse 302 CA2-IG-FC up | 0.007318 | 0.253000 | 5.52 | 27.16 |
| 8 | Human OPC L1-6 PDGFRA COL20A1 down | 0.009788 | 0.296100 | 2.32 | 10.73 |
| 9 | Mouse 377 Micro up | 0.013090 | 0.334500 | 6.45 | 27.95 |
| 10 | Mouse 358 Astro up | 0.013820 | 0.334500 | 6.31 | 27.03 |

Table S24: frontal cortex down(brain up)

| Index | Name | P-value | Adjusted p-value | Odds Ratio | Combined score |
|---|---|---|---|---|---|
| 1 | Mouse 370 Endo down | 0.00005046 | 0.01655 | 4.42 | 43.78 |
| 2 | Mouse 376 VLMC down | 0.00026610 | 0.03233 | 4.67 | 38.41 |
| 3 | Human Inh L5-6 PVALB FAM150B down | 0.00049240 | 0.03233 | 21.95 | 167.17 |
| 4 | Mouse 377 Micro down | 0.00049280 | 0.03233 | 4.76 | 36.25 |
| 5 | Mouse 378 Micro down | 0.00049280 | 0.03233 | 4.76 | 36.25 |
| 6 | Mouse 356 Astro down | 0.00104000 | 0.04918 | 3.53 | 24.26 |
| 7 | Mouse 374 VLMC down | 0.00107900 | 0.04918 | 3.81 | 26.00 |
| 8 | Human VLMC L1-5 PDGFRA COLEC12 down | 0.00119900 | 0.04918 | 3.24 | 21.80 |
| 9 | Human Exc L3-5 RORB RPRM down | 0.00158100 | 0.05762 | 40.59 | 261.80 |
| 10 | Mouse 123 DG down | 0.00200500 | 0.06458 | 13.06 | 81.16 |
| 11 | Mouse 76 Sst down | 0.00216600 | 0.06458 | 33.82 | 207.50 |
| 12 | Human Inh L3-5 VIP HS3ST3A1 down | 0.00237000 | 0.06479 | 12.28 | 74.22 |
| 13 | Human Inh L1-6 PVALB COL15A1 down | 0.00306600 | 0.07736 | 11.16 | 64.58 |
| 14 | Mouse 373 SMC up | 0.00387600 | 0.09080 | 10.23 | 56.79 |
| 15 | Human Astro L1-6 FGFR3 PLCG1 down | 0.00449100 | 0.09374 | 2.71 | 14.63 |
| 16 | Mouse 366 Oligo up | 0.00497400 | 0.09374 | 6.19 | 32.84 |
| 17 | Mouse 57 Vip down | 0.00497400 | 0.09374 | 4.82 | 25.56 |
| 18 | Mouse 379 PVM down | 0.00514400 | 0.09374 | 3.55 | 18.72 |
| 19 | Mouse 363 Oligo down | 0.00583900 | 0.09585 | 2.75 | 14.12 |
| 20 | Human Oligo L2-6 OPALIN FTH1P3 down | 0.00584500 | 0.09585 | 2.61 | 13.40 |
| 21 | Mouse 238 L5 PT CTX up | 0.00713700 | 0.11150 | 4.40 | 21.77 |
| 22 | Human Endo L2-5 NOSTRIN SRGN down | 0.00862500 | 0.12800 | 2.74 | 13.01 |
| 23 | Mouse 359 OPC down | 0.00936200 | 0.12800 | 2.55 | 11.91 |
| 24 | Mouse 373 SMC down | 0.01139000 | 0.12800 | 3.03 | 13.57 |
| 25 | Mouse 137 L2 IT ENTl up | 0.01204000 | 0.12800 | 6.66 | 29.43 |

#### $\ell_1 = 2$ only

Table S25: left cerebral cortex up1

| Index | Name | P-value | Adjusted p-value | Odds Ratio | Combined score |
|---|---|---|---|---|---|
| 1 | Mouse 125 DG down | 0.005481 | 0.6952 | 3.51 | 18.26 |
| 2 | Mouse 129 L2/3 IT APr down | 0.015530 | 0.6952 | 4.39 | 18.30 |
| 3 | Mouse 312 NP SUB up | 0.017060 | 0.6952 | 10.67 | 43.42 |
| 4 | Human Exc L3-5 RORB RPRM up | 0.024200 | 0.6952 | 3.20 | 11.91 |
| 5 | Mouse 124 DG down | 0.030360 | 0.6952 | 3.55 | 12.40 |
| 6 | Mouse 168 L2/3 IT ENTl down | 0.037420 | 0.6952 | 6.86 | 22.55 |
| 7 | Mouse 152 L2 IT ENTm down | 0.039310 | 0.6952 | 28.71 | 92.90 |
| 8 | Mouse 165 L2/3 IT ENTl down | 0.039310 | 0.6952 | 28.71 | 92.90 |
| 9 | Mouse 333 L6b/CT ENT up | 0.043170 | 0.6952 | 6.33 | 19.88 |
| 10 | Mouse 96 Sst up | 0.043180 | 0.6952 | 1.65 | 5.19 |

Table S26: left cerebral cortex down1(brain up)

| Index | Name | P-value | Adjusted p-value | Odds Ratio | Combined score |
|---|---|---|---|---|---|
| 1 | Mouse 22 Ndnf HPF down | 0.00003345 | 0.007107 | 61.52 | 633.95 |
| 2 | Mouse 33 Sncg down | 0.00004230 | 0.007107 | 23.65 | 238.16 |
| 3 | Mouse 121 DG down | 0.00010380 | 0.008897 | 5.32 | 48.83 |
| 4 | Mouse 122 DG down | 0.00010590 | 0.008897 | 5.31 | 48.58 |
| 5 | Mouse 125 DG down | 0.00028030 | 0.018210 | 4.63 | 37.89 |
| 6 | Mouse 19 Pax6 down | 0.00032510 | 0.018210 | 25.61 | 205.71 |
| 7 | Mouse 100 Sst down | 0.00059340 | 0.026600 | 20.48 | 152.19 |
| 8 | Mouse 360 OPC down | 0.00078530 | 0.026600 | 3.07 | 21.94 |
| 9 | Mouse 38 Sncg down | 0.00083290 | 0.026600 | 18.07 | 128.13 |
| 10 | Mouse 57 Vip down | 0.00083500 | 0.026600 | 5.87 | 41.62 |

Table S27: right cerebral cortex up1

| Index | Name | P-value | Adjusted p-value | Odds Ratio | Combined score |
|---|---|---|---|---|---|
| 1 | Mouse 312 NP SUB up | 0.01706 | 0.6216 | 10.67 | 43.42 |
| 2 | Mouse 70 Sst up | 0.03306 | 0.6216 | 7.36 | 25.11 |
| 3 | Mouse 152 L2 IT ENTm down | 0.03931 | 0.6216 | 28.71 | 92.90 |
| 4 | Mouse 165 L2/3 IT ENTl down | 0.03931 | 0.6216 | 28.71 | 92.90 |
| 5 | Mouse 333 L6b/CT ENT up | 0.04317 | 0.6216 | 6.33 | 19.88 |
| 6 | Mouse 148 L2 IT PAR down | 0.04412 | 0.6216 | 25.12 | 78.38 |
| 7 | Mouse 20 Ndnf HPF up | 0.04419 | 0.6216 | 3.97 | 12.37 |
| 8 | Mouse 293 CA3 down | 0.05366 | 0.6216 | 20.09 | 58.77 |
| 9 | Mouse 132 L2/3 IT PPP down | 0.05915 | 0.6216 | 3.51 | 9.91 |
| 10 | Mouse 125 DG down | 0.06211 | 0.6216 | 2.44 | 6.78 |

Table S28: right cerebral cortex down1(brain up)

| Index | Name | P-value | Adjusted p-value | Odds Ratio | Combined score |
|---|---|---|---|---|---|
| 1 | Mouse 100 Sst down | 0.00002154 | 0.006642 | 28.55 | 306.79 |
| 2 | Mouse 33 Sncg down | 0.00004230 | 0.006642 | 23.65 | 238.16 |
| 3 | Mouse 93 Sst down | 0.00011090 | 0.011610 | 38.44 | 350.03 |
| 4 | Mouse 5 Lamp5 Lhx6 down | 0.00020960 | 0.016450 | 10.12 | 85.69 |
| 5 | Mouse 119 Pvalb Vipr2 down | 0.00027950 | 0.017010 | 14.01 | 114.65 |
| 6 | Mouse 19 Pax6 down | 0.00032510 | 0.017010 | 25.61 | 205.71 |
| 7 | Mouse 56 Vip down | 0.00041990 | 0.018840 | 6.73 | 52.32 |
| 8 | Mouse 101 Sst down | 0.00063210 | 0.024810 | 11.16 | 82.23 |
| 9 | Mouse 36 Sncg down | 0.00076820 | 0.026800 | 18.62 | 133.53 |
| 10 | Mouse 71 Sst down | 0.00087090 | 0.027350 | 58.00 | 408.65 |

Table S29: hippocampal layer up1

| Index | Name | P-value | Adjusted p-value | Odds Ratio | Combined score |
|---|---|---|---|---|---|
| 1 | Mouse 96 Sst up | 0.01187 | 0.5843 | 1.88 | 8.34 |
| 2 | Mouse 132 L2/3 IT PPP down | 0.01204 | 0.5843 | 4.75 | 21.00 |
| 3 | Mouse 129 L2/3 IT APr down | 0.01553 | 0.5843 | 4.39 | 18.30 |
| 4 | Mouse 276 CA1-ProS up | 0.01871 | 0.5843 | 10.13 | 40.31 |
| 5 | Mouse 19 Pax6 up | 0.02131 | 0.5843 | 9.42 | 36.27 |
| 6 | Mouse 92 Sst up | 0.02499 | 0.5843 | 8.62 | 31.80 |
| 7 | Mouse 372 SMC down | 0.02952 | 0.5843 | 2.70 | 9.50 |
| 8 | Mouse 70 Sst up | 0.03306 | 0.5843 | 7.36 | 25.11 |
| 9 | Mouse 234 L3 RSP-ACA down | 0.03332 | 0.5843 | 2.41 | 8.19 |
| 10 | Mouse 373 SMC down | 0.03607 | 0.5843 | 2.57 | 8.53 |

Table S30: hippocampal layer down1(brain up)

| Index | Name | P-value | Adjusted p-value | Odds Ratio | Combined score |
|---|---|---|---|---|---|
| 1 | Mouse 100 Sst down | 0.00002154 | 0.007488 | 28.55 | 306.79 |
| 2 | Mouse 33 Sncg down | 0.00004230 | 0.007488 | 23.65 | 238.16 |
| 3 | Mouse 93 Sst down | 0.00011090 | 0.013080 | 38.44 | 350.03 |
| 4 | Mouse 366 Oligo down | 0.00016390 | 0.014510 | 3.65 | 31.84 |
| 5 | Mouse 5 Lamp5 Lhx6 down | 0.00020960 | 0.014840 | 10.12 | 85.69 |
| 6 | Mouse 119 Pvalb Vipr2 down | 0.00027950 | 0.016440 | 14.01 | 114.65 |
| 7 | Mouse 19 Pax6 down | 0.00032510 | 0.016440 | 25.61 | 205.71 |
| 8 | Mouse 56 Vip down | 0.00041990 | 0.018580 | 6.73 | 52.32 |
| 9 | Mouse 362 Oligo down | 0.00058200 | 0.022020 | 3.35 | 24.93 |
| 10 | Mouse 101 Sst down | 0.00063210 | 0.022020 | 11.16 | 82.23 |

Table S31: frontal cortex up1

| Index | Name | P-value | Adjusted p-value | Odds Ratio | Combined score |
|---|---|---|---|---|---|
| 1 | Mouse 118 Pvalb down | 0.0005292 | 0.05362 | 8.19 | 61.82 |
| 2 | Mouse 75 Sst down | 0.0005414 | 0.05362 | 21.19 | 159.39 |
| 3 | Mouse 86 Sst down | 0.0010850 | 0.05362 | 50.74 | 346.39 |
| 4 | Mouse 106 Pvalb down | 0.0012090 | 0.05362 | 15.75 | 105.81 |
| 5 | Mouse 117 Pvalb down | 0.0012090 | 0.05362 | 15.75 | 105.81 |
| 6 | Mouse 326 L6 CT CTX down | 0.0013220 | 0.05362 | 45.10 | 298.98 |
| 7 | Mouse 114 Pvalb down | 0.0014780 | 0.05362 | 14.62 | 95.30 |
| 8 | Mouse 129 L2/3 IT APr down | 0.0027280 | 0.08141 | 5.58 | 32.94 |
| 9 | Mouse 95 Sst down | 0.0032050 | 0.08141 | 27.05 | 155.37 |
| 10 | Mouse 116 Pvalb down | 0.0032050 | 0.08141 | 27.05 | 155.37 |

Table S32: frontal cortex down1(brain up)

| Index | Name | P-value | Adjusted p-value | Odds Ratio | Combined score |
|---|---|---|---|---|---|
| 1 | Mouse 288 CA1 down | 0.0008709 | 0.09914 | 58.00 | 408.65 |
| 2 | Mouse 276 CA1-ProS down | 0.0013220 | 0.09914 | 45.10 | 298.98 |
| 3 | Mouse 293 CA3 down | 0.0013220 | 0.09914 | 45.10 | 298.98 |
| 4 | Mouse 88 Sst up | 0.0023700 | 0.12890 | 12.28 | 74.22 |
| 5 | Mouse 90 Sst up | 0.0035380 | 0.12890 | 10.58 | 59.72 |
| 6 | Mouse 95 Sst up | 0.0040040 | 0.12890 | 23.87 | 131.77 |
| 7 | Mouse 87 Sst up | 0.0044340 | 0.12890 | 22.54 | 122.14 |
| 8 | Mouse 297 CA3 down | 0.0053560 | 0.12890 | 20.29 | 106.08 |
| 9 | Mouse 361 Oligo down | 0.0058920 | 0.12890 | 2.60 | 13.37 |
| 10 | Mouse 362 Oligo down | 0.0062040 | 0.12890 | 2.72 | 13.82 |

#### $\ell_1 = 3$ only

Table S33: left cerebral cortex up2

| Index | Name | P-value | Adjusted p-value | Odds Ratio | Combined score |
|---|---|---|---|---|---|
| 1 | Mouse 70 Sst up | 0.002918 | 0.5107 | 11.37 | 66.34 |
| 2 | Mouse 261 L6 Car3 up | 0.016270 | 0.8030 | 3.11 | 12.82 |
| 3 | Mouse 72 Sst up | 0.022840 | 0.8030 | 5.18 | 19.59 |
| 4 | Mouse 235 L3 RSP-ACA up | 0.026920 | 0.8030 | 8.27 | 29.89 |
| 5 | Mouse 302 CA2-IG-FC up | 0.041460 | 0.8030 | 4.07 | 12.96 |
| 6 | Human Inh L5-6 PVALB SST CRHR2 down | 0.044120 | 0.8030 | 25.12 | 78.38 |
| 7 | Mouse 182 L2/3 IT CTX up | 0.054270 | 0.8030 | 5.54 | 16.15 |
| 8 | Human Inh L3-5 SST CDH3 down | 0.056860 | 0.8030 | 5.39 | 15.47 |
| 9 | Mouse 293 CA3 up | 0.058200 | 0.8030 | 2.49 | 7.08 |
| 10 | Mouse 15 Lamp5 up | 0.058390 | 0.8030 | 18.26 | 51.88 |

Table S34: left cerebral cortex down2(brain up)

| Index | Name | P-value | Adjusted p-value | Odds Ratio | Combined score |
|---|---|---|---|---|---|
| 1 | Mouse 8 Lamp5 down | 0.0004032 | 0.06473 | 23.64 | 184.78 |
| 2 | Mouse 314 NP SUB down | 0.0004464 | 0.06473 | 22.76 | 175.61 |
| 3 | Mouse 300 CA3 down | 0.0018620 | 0.13710 | 36.90 | 231.95 |
| 4 | Human Inh L1 VIP KLHDC8B down | 0.0018910 | 0.13710 | 13.35 | 83.70 |
| 5 | Mouse 299 CA3 down | 0.0024910 | 0.14450 | 31.22 | 187.17 |
| 6 | Mouse 119 Pvalb Vipr2 down | 0.0038760 | 0.17710 | 10.23 | 56.79 |
| 7 | Mouse 332 L6 CT ENTm down | 0.0048850 | 0.17710 | 21.35 | 113.64 |
| 8 | Mouse 10 Lamp5 down | 0.0048850 | 0.17710 | 21.35 | 113.64 |
| 9 | Mouse 296 CA3 down | 0.0085960 | 0.25790 | 15.60 | 74.20 |
| 10 | Mouse 46 Vip down | 0.0089080 | 0.25790 | 7.47 | 35.29 |
| 11 | Mouse 121 DG down | 0.0109700 | 0.25790 | 3.41 | 15.37 |
| 12 | Mouse 110 Pvalb down | 0.0118100 | 0.25790 | 13.08 | 58.06 |
| 13 | Human Inh L1 LAMP5 RAB11FIP1 down | 0.0125100 | 0.25790 | 12.67 | 55.51 |
| 14 | Human Inh L6 SST TH down | 0.0139600 | 0.25790 | 11.92 | 50.93 |
| 15 | Mouse 112 Pvalb down | 0.0147100 | 0.25790 | 11.58 | 48.87 |
| 16 | Mouse 56 Vip down | 0.0160700 | 0.25790 | 4.35 | 17.95 |
| 17 | Mouse 33 Sncg down | 0.0162600 | 0.25790 | 10.96 | 45.13 |
| 18 | Human Inh L5-6 LAMP5 CRABP1 down | 0.0170600 | 0.25790 | 10.67 | 43.42 |
| 19 | Mouse 106 Pvalb down | 0.0187100 | 0.25790 | 10.13 | 40.31 |
| 20 | Mouse 3 Lamp5 Lhx6 down | 0.0187100 | 0.25790 | 10.13 | 40.31 |
| 21 | Mouse 353 NP PPP down | 0.0195600 | 0.25790 | 9.89 | 38.89 |
| 22 | Mouse 103 Sst down | 0.0213100 | 0.25790 | 9.42 | 36.27 |
| 23 | Mouse 114 Pvalb down | 0.0213100 | 0.25790 | 9.42 | 36.27 |
| 24 | Human Inh L2-5 VIP SOX11 up | 0.0218700 | 0.25790 | 5.27 | 20.16 |
| 25 | Mouse 302 CA2-IG-FC down | 0.0231200 | 0.25790 | 9.00 | 33.92 |

Table S35: right cerebral cortex up2

| Index | Name | P-value | Adjusted p-value | Odds Ratio | Combined score |
|---|---|---|---|---|---|
| 1 | Mouse 375 VLMC down | 0.05182 | 0.8934 | 2.34 | 6.92 |
| 2 | Mouse 261 L6 Car3 up | 0.05307 | 0.8934 | 2.56 | 7.51 |
| 3 | Human Inh L5-6 SST PIK3CD down | 0.06082 | 0.8934 | 5.19 | 14.52 |
| 4 | Mouse 89 Sst up | 0.06082 | 0.8934 | 5.19 | 14.52 |
| 5 | Mouse 28 Sncg up | 0.06625 | 0.8934 | 2.72 | 7.39 |
| 6 | Mouse 330 L6 CT CTX down | 0.07710 | 0.8934 | 13.39 | 34.32 |
| 7 | Mouse 238 L5 PT CTX down | 0.08171 | 0.8934 | 12.55 | 31.44 |
| 8 | Mouse 142 L3 IT ENTm down | 0.09088 | 0.8934 | 11.16 | 26.76 |
| 9 | Mouse 259 L6 Car3 up | 0.09996 | 0.8934 | 10.04 | 23.12 |
| 10 | Mouse 308 L5 NP CTX down | 0.10450 | 0.8934 | 9.56 | 21.60 |

Table S36: right cerebral cortex down2(brain up)

| Index | Name | P-value | Adjusted p-value | Odds Ratio | Combined score |
|---|---|---|---|---|---|
| 1 | Mouse 10 Lamp5 down | 0.0001511 | 0.03598 | 34.16 | 300.54 |
| 2 | Mouse 8 Lamp5 down | 0.0004032 | 0.03598 | 23.64 | 184.78 |
| 3 | Mouse 314 NP SUB down | 0.0004464 | 0.03598 | 22.76 | 175.61 |
| 4 | Mouse 118 Pvalb down | 0.0005292 | 0.03598 | 8.19 | 61.82 |
| 5 | Mouse 144 L3 IT ENTl down | 0.0010850 | 0.05479 | 50.74 | 346.39 |
| 6 | Mouse 3 Lamp5 Lhx6 down | 0.0012090 | 0.05479 | 15.75 | 105.81 |
| 7 | Mouse 300 CA3 down | 0.0018620 | 0.07237 | 36.90 | 231.95 |
| 8 | Mouse 140 L3 IT ENTm down | 0.0021660 | 0.07364 | 33.82 | 207.50 |
| 9 | Mouse 299 CA3 down | 0.0024910 | 0.07528 | 31.22 | 187.17 |
| 10 | Mouse 56 Vip down | 0.0028530 | 0.07761 | 5.52 | 32.33 |

Table S37: hippocampal layer up2

| Index | Name | P-value | Adjusted p-value | Odds Ratio | Combined score |
|---|---|---|---|---|---|
| 1 | Mouse 259 L6 Car3 up | 0.004885 | 0.7768 | 21.35 | 113.64 |
| 2 | Mouse 9 Lamp5 up | 0.023120 | 0.8555 | 9.00 | 33.92 |
| 3 | Mouse 374 VLMC down | 0.043830 | 0.8555 | 2.44 | 7.64 |
| 4 | Mouse 10 Lamp5 up | 0.051720 | 0.8555 | 5.70 | 16.88 |
| 5 | Mouse 229 L6 IT CTX up | 0.052560 | 0.8555 | 2.95 | 8.70 |
| 6 | Mouse 231 L6 IT CTX up | 0.052990 | 0.8555 | 5.62 | 16.51 |
| 7 | Mouse 261 L6 Car3 up | 0.053070 | 0.8555 | 2.56 | 7.51 |
| 8 | Mouse 28 Sncg up | 0.066250 | 0.8555 | 2.72 | 7.39 |
| 9 | Mouse 55 Vip up | 0.067790 | 0.8555 | 15.45 | 41.59 |
| 10 | Human Exc L3 RORB OTOGL up | 0.077100 | 0.8555 | 13.39 | 34.32 |
| 11 | Mouse 262 L6 Car3 up | 0.079830 | 0.8555 | 3.08 | 7.78 |
| 12 | Human Exc L3-5 RORB RPRM up | 0.081620 | 0.8555 | 2.53 | 6.33 |
| 13 | Mouse 355 V3d up | 0.081620 | 0.8555 | 2.53 | 6.33 |
| 14 | Mouse 370 Endo down | 0.083840 | 0.8555 | 2.05 | 5.08 |
| 15 | Mouse 373 SMC down | 0.098590 | 0.8555 | 2.11 | 4.89 |
| 16 | Mouse 368 Oligo down | 0.103200 | 0.8555 | 1.93 | 4.38 |
| 17 | Mouse 329 L6 CT CTX up | 0.104500 | 0.8555 | 9.56 | 21.60 |
| 18 | Human Exc L5 RORB MED8 up | 0.131000 | 0.8555 | 7.43 | 15.11 |
| 19 | Mouse 8 Lamp5 up | 0.139700 | 0.8555 | 6.92 | 13.62 |
| 20 | Mouse 379 PVM down | 0.159100 | 0.8555 | 1.95 | 3.59 |
| 21 | Mouse 30 Sncg up | 0.159500 | 0.8555 | 1.69 | 3.10 |
| 22 | Mouse 233 L6 IT ENTl up | 0.161000 | 0.8555 | 5.90 | 10.78 |
| 23 | Mouse 118 Pvalb up | 0.165200 | 0.8555 | 5.73 | 10.32 |
| 24 | Mouse 60 Sst Chodl up | 0.173600 | 0.8555 | 5.42 | 9.50 |
| 25 | Human Inh L1-6 LAMP5 CA1 up | 0.181800 | 0.8555 | 5.14 | 8.77 |

Table S38: hippocampal layer down2(brain up)

| Index | Name | P-value | Adjusted p-value | Odds Ratio | Combined score |
|---|---|---|---|---|---|
| 1 | Mouse 314 NP SUB down | 0.0004464 | 0.06138 | 22.76 | 175.61 |
| 2 | Mouse 118 Pvalb down | 0.0005292 | 0.06138 | 8.19 | 61.82 |
| 3 | Mouse 306 L5 NP CTX down | 0.0016770 | 0.12970 | 13.96 | 89.20 |
| 4 | Mouse 56 Vip down | 0.0028530 | 0.16490 | 5.52 | 32.33 |
| 5 | Mouse 142 L3 IT ENTm down | 0.0040040 | 0.16490 | 23.87 | 131.77 |
| 6 | Mouse 10 Lamp5 down | 0.0048850 | 0.16490 | 21.35 | 113.64 |
| 7 | Mouse 57 Vip down | 0.0049740 | 0.16490 | 4.82 | 25.56 |
| 8 | Mouse 121 DG down | 0.0109700 | 0.28630 | 3.41 | 15.37 |
| 9 | Mouse 122 DG down | 0.0111100 | 0.28630 | 3.40 | 15.29 |
| 10 | Mouse 120 DG down | 0.0235700 | 0.35620 | 2.85 | 10.68 |

Table S39: frontal cortex up2

| Index | Name | P-value | Adjusted p-value | Odds Ratio | Combined score |
|---|---|---|---|---|---|
| 1 | Mouse 259 L6 Car3 up | 0.004885 | 0.6287 | 21.35 | 113.64 |
| 2 | Mouse 31 Sncg down | 0.010470 | 0.6287 | 13.98 | 63.75 |
| 3 | Mouse 20 Ndnf HPF down | 0.011130 | 0.6287 | 13.52 | 60.80 |
| 4 | Mouse 373 SMC down | 0.036070 | 0.6287 | 2.57 | 8.53 |
| 5 | Mouse 141 L3 IT ENTm up | 0.039310 | 0.6287 | 28.71 | 92.90 |
| 6 | Mouse 261 L6 Car3 up | 0.053070 | 0.6287 | 2.56 | 7.51 |
| 7 | Mouse 376 VLMC down | 0.060630 | 0.6287 | 2.46 | 6.89 |
| 8 | Human Oligo L2-6 OPALIN FTH1P3 up | 0.062160 | 0.6287 | 5.12 | 14.22 |
| 9 | Mouse 308 L5 NP CTX up | 0.067790 | 0.6287 | 15.45 | 41.59 |
| 10 | Mouse 310 L5 NP CTX up | 0.069020 | 0.6287 | 4.81 | 12.87 |
| 11 | Mouse 142 L3 IT ENTm up | 0.072460 | 0.6287 | 14.35 | 37.66 |
| 12 | Human Exc L3 RORB OTOGL up | 0.077100 | 0.6287 | 13.39 | 34.32 |
| 13 | Mouse 351 L6b CTX down | 0.077100 | 0.6287 | 13.39 | 34.32 |
| 14 | Mouse 262 L6 Car3 up | 0.079830 | 0.6287 | 3.08 | 7.78 |
| 15 | Mouse 15 Lamp5 down | 0.081710 | 0.6287 | 12.55 | 31.44 |
| 16 | Mouse 37 Sncg down | 0.086310 | 0.6287 | 11.81 | 28.94 |
| 17 | Mouse 158 L2/3 IT ENTl up | 0.090880 | 0.6287 | 11.16 | 26.76 |
| 18 | Mouse 185 L2 IT RSP-ACA down | 0.097110 | 0.6287 | 2.82 | 6.57 |
| 19 | Mouse 329 L6 CT CTX up | 0.104500 | 0.6287 | 9.56 | 21.60 |
| 20 | Mouse 140 L3 IT ENTm up | 0.117800 | 0.6287 | 8.37 | 17.89 |
| 21 | Mouse 258 L6 Car3 down | 0.131000 | 0.6287 | 7.43 | 15.11 |
| 22 | Mouse 30 Sncg down | 0.131000 | 0.6287 | 7.43 | 15.11 |
| 23 | Mouse 8 Lamp5 down | 0.135400 | 0.6287 | 7.17 | 14.34 |
| 24 | Mouse 97 Sst up | 0.137600 | 0.6287 | 1.58 | 3.13 |
| 25 | Human Inh L3-6 VIP UG0898H09 down | 0.139700 | 0.6287 | 6.92 | 13.62 |

Table S40: frontal cortex down2(brain up)

| Index | Name | P-value | Adjusted p-value | Odds Ratio | Combined score |
|---|---|---|---|---|---|
| 1 | Mouse 112 Pvalb up | 0.0003551 | 0.06854 | 8.98 | 71.30 |
| 2 | Mouse 110 Pvalb up | 0.0043500 | 0.31430 | 3.07 | 16.67 |
| 3 | Mouse 116 Pvalb up | 0.0048850 | 0.31430 | 21.35 | 113.64 |
| 4 | Mouse 113 Pvalb up | 0.0109300 | 0.38000 | 4.89 | 22.10 |
| 5 | Mouse 111 Pvalb up | 0.0110400 | 0.38000 | 3.05 | 13.75 |
| 6 | Mouse 115 Pvalb up | 0.0118100 | 0.38000 | 13.08 | 58.06 |
| 7 | Mouse 109 Pvalb up | 0.0142200 | 0.39200 | 4.51 | 19.20 |
| 8 | Mouse 355 V3d up | 0.0242000 | 0.45970 | 3.20 | 11.91 |
| 9 | Mouse 57 Vip down | 0.0246200 | 0.45970 | 3.80 | 14.06 |
| 10 | Mouse 105 Pvalb up | 0.0259500 | 0.45970 | 8.44 | 30.82 |
| 11 | Mouse 114 Pvalb up | 0.0274900 | 0.45970 | 4.82 | 17.31 |
| 12 | Mouse 118 Pvalb down | 0.0285800 | 0.45970 | 4.74 | 16.85 |
| 13 | Mouse 70 Sst up | 0.0330600 | 0.49080 | 7.36 | 25.11 |
| 14 | Mouse 80 Sst up | 0.0467600 | 0.57230 | 6.04 | 18.50 |
| 15 | Mouse 10 Lamp5 up | 0.0517200 | 0.57230 | 5.70 | 16.88 |
| 16 | Mouse 334 L6b/CT ENT down | 0.0536600 | 0.57230 | 20.09 | 58.77 |
| 17 | Human Inh L3-5 SST CDH3 down | 0.0568600 | 0.57230 | 5.39 | 15.47 |
| 18 | Mouse 98 Sst up | 0.0632200 | 0.57230 | 3.41 | 9.41 |
| 19 | Human Exc L6 THEMIS SLN up | 0.0657300 | 0.57230 | 3.35 | 9.12 |
| 20 | Human Oligo L5-6 OPALIN LDLRAP1 up | 0.0688000 | 0.57230 | 2.69 | 7.19 |
| 21 | Human Exc L6 THEMIS LINC00343 up | 0.0690200 | 0.57230 | 4.81 | 12.87 |
| 22 | Mouse 97 Sst up | 0.0718300 | 0.57230 | 1.77 | 4.67 |
| 23 | Mouse 142 L3 IT ENTm up | 0.0724600 | 0.57230 | 14.35 | 37.66 |
| 24 | Mouse 56 Vip down | 0.0726200 | 0.57230 | 3.21 | 8.41 |
| 25 | Mouse 73 Sst up | 0.0771000 | 0.57230 | 13.39 | 34.32 |
