## Supplementary figures and images for "Somatostatin-expressing neurons regulate sleep deprivation and recovery"

### Figure_S1.pdf

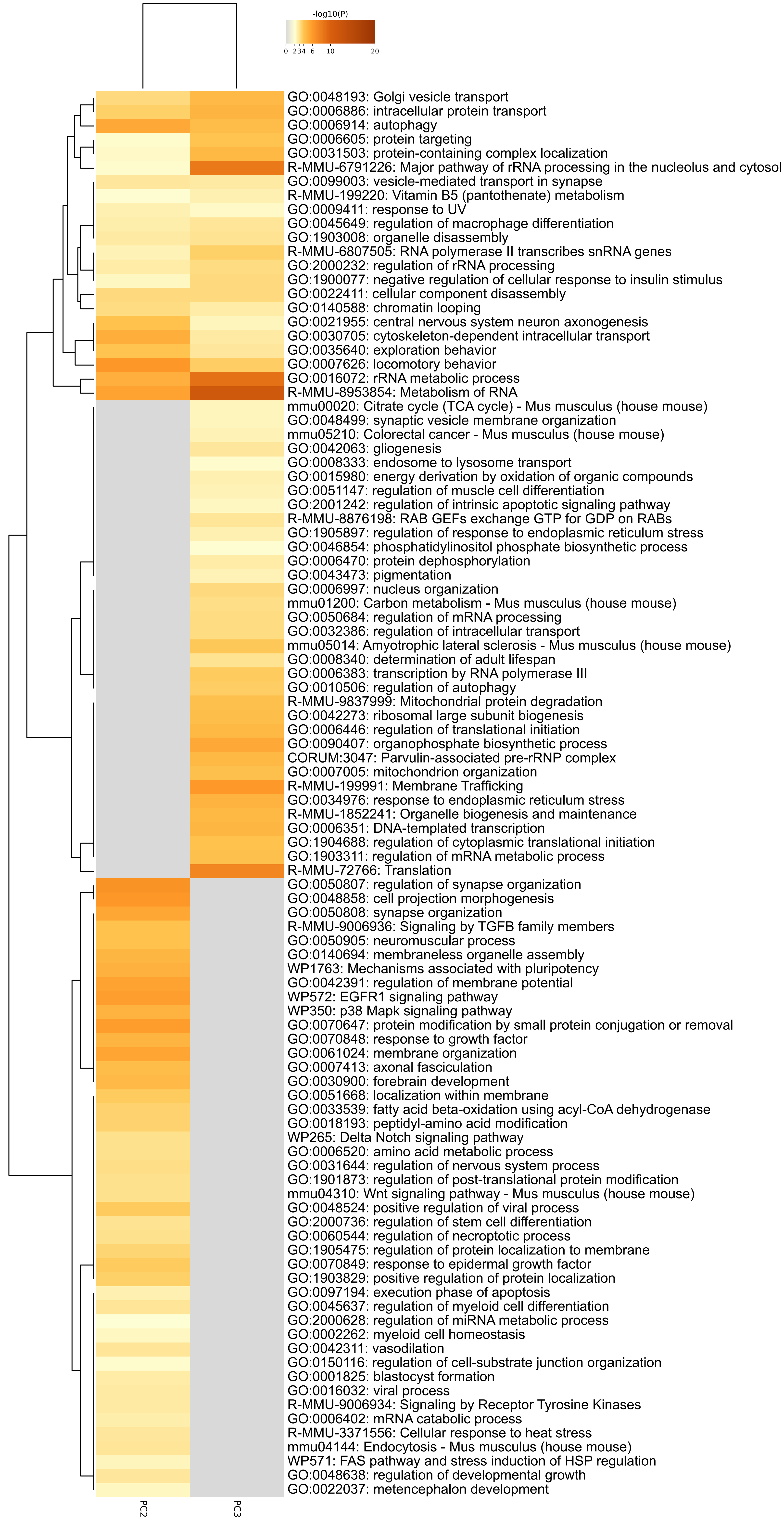
